# A conserved pituitary response to tissue loss controls regeneration across vertebrates

**DOI:** 10.64898/2026.10.31.762810

**Authors:** Noah Lopez, Julia C. Paoli, Duygu Payzin-Dogru, Hani Singer, Tim Froitzheim, Luciana Landívar-Uriona, Antzela Zgouro, Emily de Rezende, Ava R. Moss, S.Y. Celeste Wu, Bradley K. Fox, Reilly S. Merlo, Ryan J.A. Chang, Tyler R. Goodearly, Laurel Shugart, Ashley Juarez, Erica Chen, Anthony E. Striker, Anna Luong, Ryan T. Kim, Miranda X. Thai, Anne T. Harrington, Tia S. Patel, Matthew Chen, Anne S. Golding, Michael J. Raymond, Jessica A. Lehoczky, Andre P. Seale, Jason P. Breves, Jessica L. Whited

## Abstract

Regenerating complex tissues requires coordination between local cellular responses and body-wide systemic signals. As a central endocrine regulator, the pituitary is well positioned to coordinate this process. Here, we identify the pituitary as an injury-responsive regulator of axolotl limb and tilapia fin regeneration. Limb loss triggered sustained transcriptional responses in the axolotl pituitary. Single-cell transcriptomics linked pituitary loss to altered gene expression in prolactin- and growth hormone-responsive limb cells. Local disruption of the prolactin receptor impaired digit formation, whereas combined hormone treatment partially restored regeneration in hypophysectomized axolotls. In tilapia, pituitary removal impaired fin regeneration and prolactin-related signaling. Mouse digits differed from axolotls and tilapia in how and when they received pituitary signals during regeneration, suggesting pituitary-dependent regrowth is evolutionarily divergent among highly regenerative vertebrates.

## Introduction

Successful appendage regeneration relies on many aspects of an animal’s biology, including the coordinated cell behaviors of exact signaling timing, successful signal responses, and correct communication between local and distant tissues. Here, we investigate how pituitary-derived endocrine signals regulate local cellular responses during appendage regeneration. The pituitary gland is ideally situated to integrate distant injury signals and shape organismal responses. Environmental and endogenous perturbation can stimulate the pituitary directly, or indirectly via hypothalamic and systemic inputs (*1–3*). These stimuli trigger the release of hormones from the pituitary that enter the circulatory system to affect physiological processes throughout the organism (*4*). While historical endocrine studies have firmly established the importance of pituitary hormones in coordinating growth, metabolism, and the maintenance of specific peripheral tissues (*5–7*), their roles in appendage regeneration remain unresolved.

Early studies identified the pituitary as a modulator of salamander limb regeneration (*8, 9*). However, these observations predated the molecular and mechanistic techniques needed to unravel how limb cells respond to pituitary signals. Moreover, there are considerable species-dependent discrepancies concerning whether the pituitary is required for limb regeneration. For example, pituitary removal (hypophysectomy) in larval *Hynobius retardatu*s (*10*) and *Ambystoma opacum* (*11*) did not block limb regeneration, while adult newts (*Triturus viridescens*) required the pituitary to initiate limb regeneration (*12*). Other reports claim that if fed well, hypophysectomized adult newts regenerate limbs, albeit more slowly (*13, 14*). Understanding how the pituitary influences regenerative success across vertebrate groups could yield crucial insights into the evolution of regeneration. Furthermore, this work may inform how pituitary signals might be harnessed for regenerative therapies. We therefore sought to use an unbiased strategy in a genetically tractable salamander model to identify which hormones the pituitary produces in response to amputation, which specific limb cell populations receive pituitary-derived signals, and how these cell types alter their biology in response.

In this work, we asked whether the pituitary functions as an injury-responsive regulator of regeneration and if this function is conserved across vertebrates. Using the axolotl (*Ambystoma mexicanum*) as our primary model, we examined how hypophysectomy affects limb regeneration. Utilizing bulk RNA-sequencing of pituitaries during normal regeneration, we identified specific temporal transcriptional programs following limb amputation. We then asked how loss of pituitary input reshapes the cellular and molecular composition of distant, regenerating tissues. Using single-cell transcriptomics, we implicated epithelial cells and fibroblasts in the regenerating limb as the cells most affected by loss of pituitary input. Further exploration of the contributions of prolactin (PRL) and its receptor (PRLR) revealed distinct deficits in blastema contribution and digit formation. To test whether pituitary-based regulation of regeneration is conserved across evolutionary time, rather than being unique to salamanders, we extended our approach to two additional species, tilapia (*Oreochromis mossambicus*) and mouse (*Mus musculus*), which diverged from the axolotl over 400 million and 350 million years ago, respectively. Our work supports a model in which the pituitary promotes the growth of the blastema which is necessary for tissue replacement in highly-regenerative species, while the deployment of specific pituitary signaling pathways after injury differs in mammals.

## Results

### Pituitary removal suppresses blastema growth

We surgically removed the pituitary from axolotls (hypophysectomy, HPS) and confirmed successful removal by loss of PRL1 protein and transcripts in the brain and adjacent structures by western blot and HCR-FISH (hybridization chain reaction fluorescent *in situ* hybridization) (**Fig**. 1A, B, C, C’). After hypophysectomy and subsequent amputation, HPS and sham-operated animals formed blastemas of similar size at 5 days post-amputation (dpa). However, by 7 and 14 dpa, HPS blastemas were significantly smaller than sham controls (**Fig**. 1D, E). EdU incorporation revealed no change in proliferation between sham and HPS blastema at 14 dpa, despite size differences (**Fig**. 1F, F’). The pituitary did not regenerate after HPS in animals up to 70 days post-surgery (**Fig**. S1).

**Figure 1.**
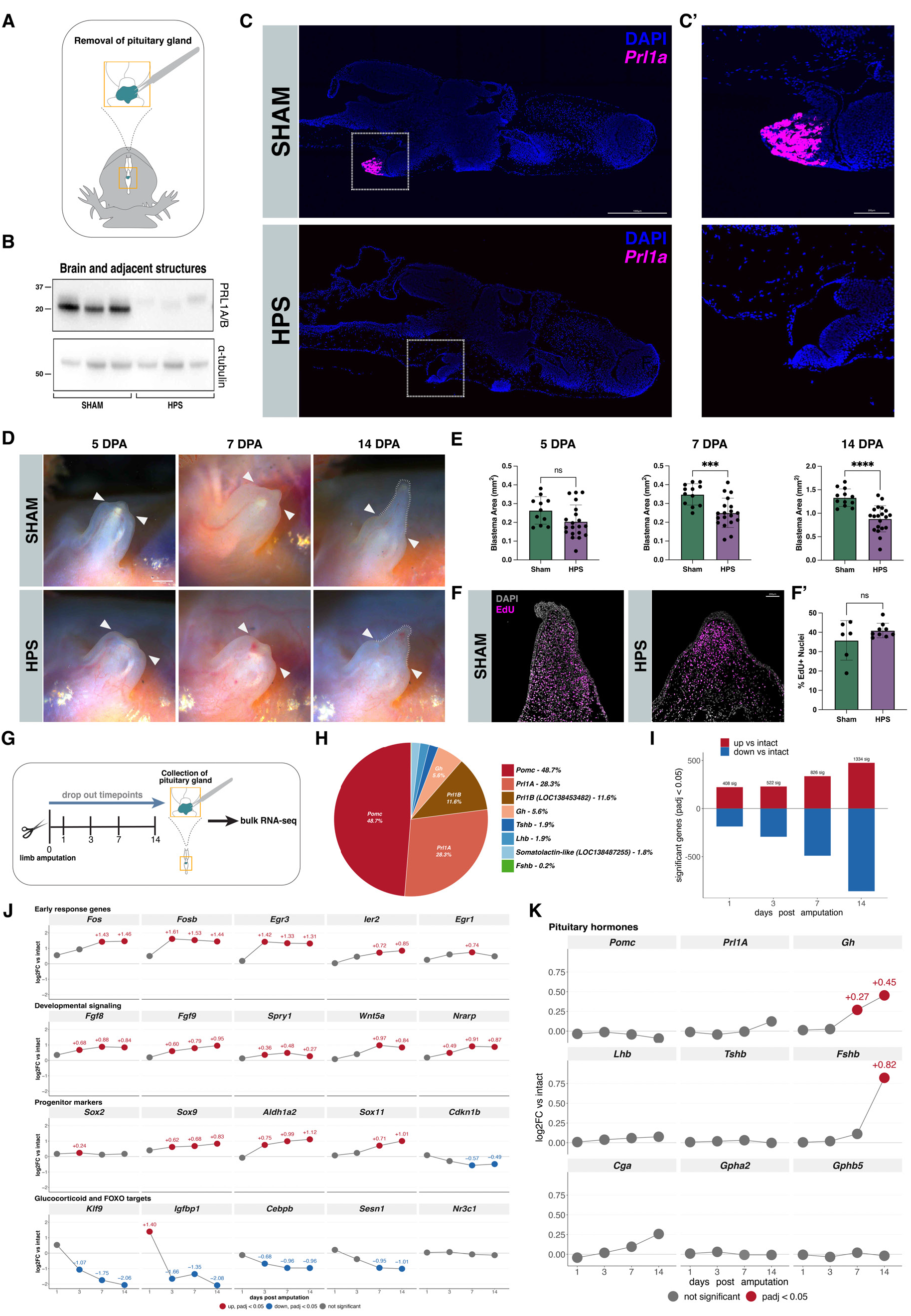
The pituitary promotes long-range growth effects on the blastema during axolotl limb regeneration. **(A)** Hypophysectomy (HPS) procedure, with the pituitary gland removed through a cranial window in the roof of the mouth. **(B)** Western blot confirming pituitary removal: brain and adjacent tissue from sham-control and hypophysectomized animals, probed for abundant pituitary hormone (PRL1) and alpha-tubulin as a loading control. **(C)** Confirmation of pituitary removal by hybridization chain reaction fluorescence in situ hybridization (HCR-FISH): *Prl1a* probe and DAPI in sagittal brain sections, Sham vs. HPS. Scale bar, 1000 µm. The boxed region is shown at higher magnification in (C′). **(C′)** Higher-magnification view of the boxed pituitary region in (C); *Prl1a* probe and DAPI. Scale bar, 200 µm **(D)** Representative Sham and HPS blastemas at 5, 7, and 14 days post-amputation (dpa); arrowheads mark the amputation plane and the dotted outline marks the blastema. Hypophysectomized axolotls show visibly impaired regeneration relative to sham controls by 14 dpa. Scale bar, 500 µm. **(E)** Blastema area (mm²) at 5 dpa, Sham vs. HPS; not significantly different. Blastema area (mm²) at 7 dpa, Sham vs. HPS; HPS significantly reduced blastema area (p < 0.001). Blastema area (mm²) at 14 dpa, Sham vs. HPS; HPS significantly reduced blastema area (p < 0.0001). **(F)** EdU labeling showing proliferation in Sham vs. HPS 14 dpa blastemas. Scale bar, 200 µm. **(F’)** EdU incorporation revealed no change in proliferation between sham and HPS 14 dpa blastema by %EdU+ nuclei. **(G)** Experimental design for bulk pituitary RNA-seq: limbs were amputated at day 0, and pituitaries were collected from intact animals and at 1, 3, 7, and 14 days post-amputation. **(H)** Share of sequenced hormone transcripts (TPM) in the intact pituitary gland. *Pomc* accounts for 48.7% and *Prl1A* for 28.3% of hormone transcripts, followed by *Prl1B* (11.6%), *Gh* (5.6%), *Tshb* and *Lhb* (1.9% each), *Somatolactin-like* (LOC138487255) (1.8%), and *Fshb* (0.2%). This panel shows the relative composition of the hormone pool. **(I)** Number of significant genes (padj < 0.05, DESeq2 Wald, BH-corrected; n = 3 samples per timepoint each pooling 5 pituitary glands) versus intact, by direction, at each timepoint after limb amputation. Significant genes increase with time, from 408 at 1 dpa to 522 at 3 dpa, 826 at 7 dpa, and 1,334 at 14 dpa. **(J)** Log2 fold change vs. intact (unshrunken) for four gene programs. Colored points are padj < 0.05. Early response genes *Fosb* and *Egr3* rose by 3 dpa, and *Fos* rose by 7 dpa. They remained elevated through 14 dpa. Developmental signaling-associated genes, *Fgf8*, *Fgf9*, and *Wnt5a*, rose by 3dpa. Progenitor markers, *Sox2*, *Sox9*, and *Aldh1a2*, rose. Glucocorticoid and FOXO targets including *Klf9*, *Cebpb*, and *Sesn1*, decreased. **(K)** Pituitary hormone transcript expression after amputation. Pituitary hormone gene expression (log2FC vs. intact) at 1, 3, 7, and 14 days post-amputation, for nine hormone and glycoprotein-subunit genes shown. Only *Gh* and *Fshb* change significantly (red): *Gh* rises at 7 and 14 dpa (+0.27, +0.45) and *Fshb* rises at 14 dpa (+0.82). Fold changes on this panel are ashr-shrunken for display, and adjusted p-values are sourced from the unshrunken fit: *Prl1A* does not change (+0.12 at 14 dpa, padj = 0.25), and *Pomc*, *Lhb*, *Tshb*, *Cga*, *Gpha2*, and *Gphb5* do not change significantly (grey).

### The pituitary mounts a broad transcriptional response to limb injury

After examining how hypophysectomy affects early limb regeneration, we investigated if the pituitary itself responds transcriptionally to a distant injury. We performed bulk RNA sequencing on pituitary glands collected at homeostasis (intact), 1, 3, 7, and 14 days following limb amputation (**Fig**. 1G). Using the intact condition, we discerned the composition of pituitary hormone transcripts. By transcript output, the gland’s hormone production is dominated by *Pomc* (48.7% of total recovered hormone transcripts), followed by *Prl1a* (28.3%), *Prl1b* (11.6%), *Gh* (5.6%), *Tshb* (1.9%), *Lhb* (1.9%), *somatolactin*-like (1.8%), and *Fshb* (0.2%) (**Fig**. 1H). Following limb amputation, the pituitary showed a clear injury response that grew in magnitude over time. Compared to the intact condition, we identified 408, 522, 826, and 1,334 differentially expressed genes (DEGs) at 1, 3, 7, and 14 dpa, respectively. (**Fig**. 1I, **Fig.** S2A). Gene set testing uncovered upregulation of pituitary gland development and endocrine system pathways and downregulation of collagen metabolic processes. Of 4,220 gene sets tested, 166 were significantly altered at 14 dpa, predominantly downward (**Fig**. S2B). Principal component analysis (PCA) visualization of samples showed clustering of amputated samples progressively farther away from the intact condition (**Fig.** S2C). This trend aligns with the temporal window in which we observed prominent blastema defects in hypophysectomized axolotls. We identified four coordinated programs of interest: **early response genes** such as *Fosb* and *Egr3* rose by 3 dpa and *Fos* by 7 dpa and remained elevated through 14 dpa, **developmental signaling**-associated genes including *Fgf8*, *Fgf9*, and *Wnt5a* rose as early as 3 dpa, **progenitor markers** associated with adult pituitary stem cells such as *Sox2*, *Sox9*, and *Aldh1a2* also rose (*15*). **Glucocorticoid and FOXO targets** including *Klf9*, *Cebpb*, and *Sesn1* fell, although the receptor *Nr3c1* itself was unchanged (**Fig**. 1J).

When we examined which hormone-encoding transcripts changed across pituitary hormone genes, only a subset showed significant injury-induced changes (**Fig**. 1K). Notably, neither *Prl1a* nor *Prl1b* transcription increased after amputation, indicating that PRL signaling is not contributing to regeneration through a simple increase in ligand transcription from the gland. However, other loci, including *Gh* and *Fshb,* showed significant, progressively increasing expression levels over the injury time course. This asymmetry is consistent with prior work showing that growth hormone-releasing hormone (*Ghrh)* increases in the hypothalamus after injury, but prolactin-releasing hormone (*Prlh)* does not (*16*). This likely reflects the different mechanisms governing GH versus PRL release. GH secretion relies more heavily on a stimulatory releasing hormone. However, hypothalamic dopamine inhibits PRL secretion and therefore requires withdrawal of dopaminergic inhibition (*17, 18*). Despite no changes in transcription, this does not rule out that the pituitary is instead modulating PRL secretion at the protein level. Together, these results indicate that regeneration is accompanied by a distinct shift in the pituitary’s transcriptional profile following limb amputation, including direct modulation of pituitary hormone production.

### Pituitary removal results in perturbation of Prlr and Ghr signaling in the limb

To characterize the shift toward pituitary dependency at the cellular level, we performed single-cell RNA sequencing (scRNA-seq) on 7 dpa regenerating limbs (including the wound epidermis, the blastema, and the underlying stump) from sham and HPS-operated animals (**Fig**. 2A). Clustering identified the expected diversity of cell types found in the blastema and adjacent tissues (epithelial cells, fibroblasts, erythroid populations, macrophages, muscle, etc.) (**Fig**. 2B, S3A). Moreover, this cell-type structure was broadly preserved across conditions (**Fig**. 2C). Pseudo-bulk differential gene expression analyses within each cell type revealed that epithelial cells and fibroblasts accounted for a large majority of DEGs between sham and HPS (**Fig**. 2D, S4A,B). We therefore focused on investigating the genes that are differentially expressed in epithelial cells and fibroblasts under the HPS condition (**Fig.** 2E, F).

**Figure 2.**
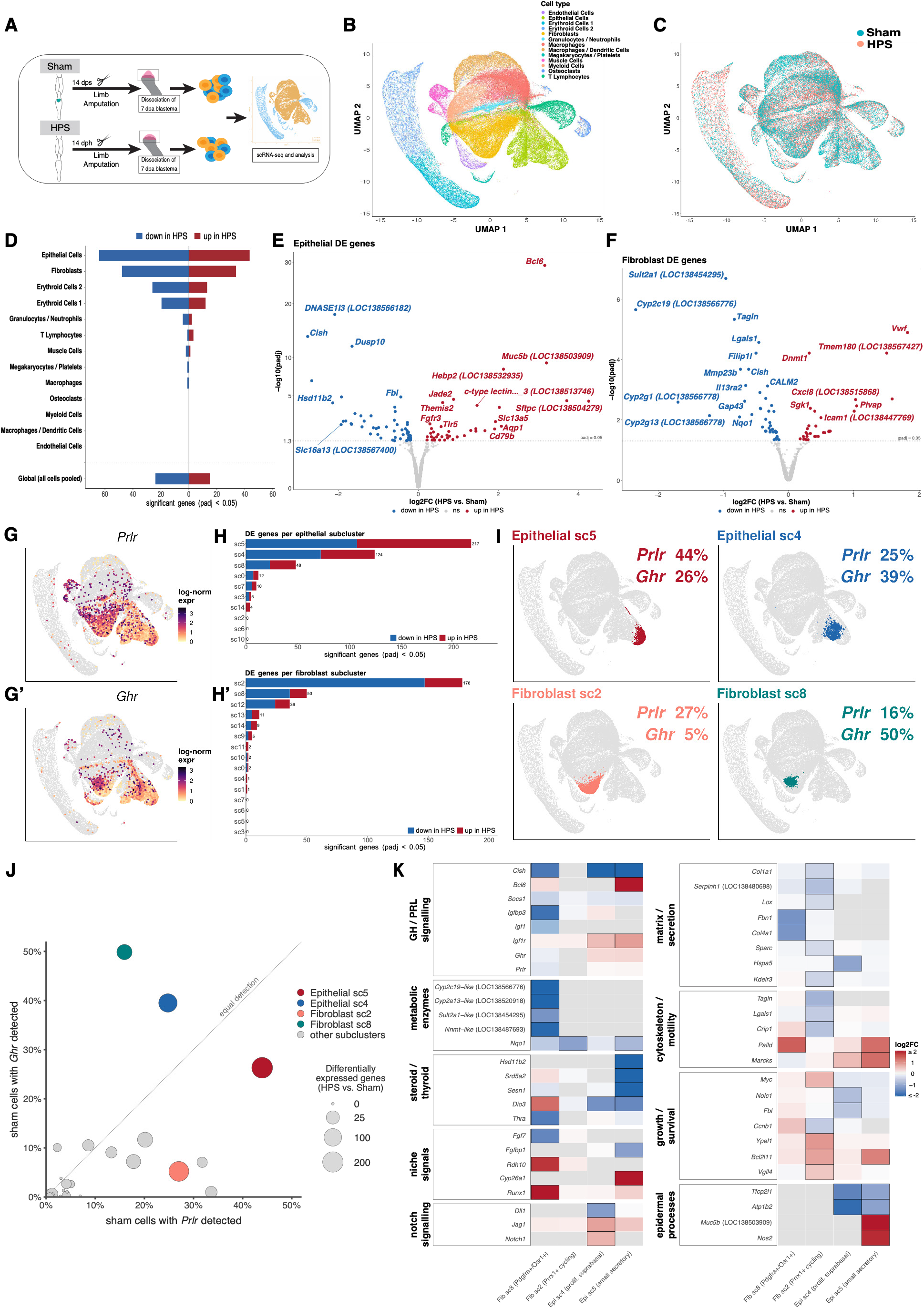
Pituitary hormone-related pathways show differential expression in axolotl limbs after hypophysectomy and amputation, and the pituitary itself responds to amputation. **(A)** Single-cell RNA-sequencing workflow: Sham and HPS limbs were amputated 14 days after the respective surgery (14 dps / 14 dph), and 7 dpa blastemas were dissociated for scRNA-seq and analysis. (B) UMAP of 146,092 cells in the 13 cell types tested for differential expression, colored by cell type.(C) UMAP of (B), colored by surgical condition (Sham vs. HPS); every cell type is represented in both conditions. (D) Number of significant pseudobulk DE genes (padj < 0.05, HPS vs. Sham) per cell type, split by direction, plus all cells pooled as a global test. Epithelial Cells (99 genes) and Fibroblasts (75 genes) carry the most significant genes, followed by Erythroid Cells 2 and 1 (36 and 29 genes, respectively). (E) Volcano plot of pseudobulk differential expression for Epithelial Cells (HPS vs. Sham; 17,433 genes tested, 99 significant at padj < 0.05, represented by the dotted line). (F) Volcano plot of pseudobulk differential expression for Fibroblasts (HPS vs. Sham; 13,752 genes tested, 75 significant at padj < 0.05, represented by the dotted line). (G) UMAP feature plots (147,253 cells) colored by log-normalized expression of *Prlr* (top) and *Ghr* (bottom). Grey = no UMI detected. *Prlr* and *Ghr* are broadly detected (3.2% and 2.3% of all cells, respectively). **(H)** Number of significant genes (padj < 0.05, HPS vs. sham; BH-corrected within subcluster) per epithelial subcluster; counted only where ≥ 3 libraries per condition carry ≥ 10 cells. The response is concentrated in two epithelial subclusters, sc5 (217 genes) and sc4 (124 genes). **(H’)** Number of significant genes per fibroblast subcluster, same design as (H). The response is concentrated in sc2 (178 genes), followed by sc8 (50 genes). **(I)** Percentage of cells expressing *Prlr* and *Ghr* in the four subclusters with the most differentially expressed genes: Epithelial sc5 (*Prlr* 44%, *Ghr* 26%), Epithelial sc4 (*Prlr* 25%, *Ghr* 39%), Fibroblast sc2 (*Prlr* 27%, *Ghr* 5%), and Fibroblast sc8 (*Prlr* 16%, *Ghr* 50%). **(J)** Scatter plot of epithelial and fibroblast subclusters (26 of 31 with ≥100 sham cells), positioned by the percentage of sham cells in which *Prlr* (x-axis) and *Ghr* (y-axis) were detected at ≥ 1 UMI, with point area proportional to the number of genes differentially expressed after hypophysectomy in that subcluster. The top two DE clusters in each cell type are colored, and the remaining 22 are left grey. Epithelial subcluster 5 (44.0% *Prlr*, 26.3% *Ghr*; 217 genes) and fibroblast subcluster 2 (27.0% *Prlr*, 5.2% *Ghr*; 178 genes) are *Prlr*-biased, while epithelial subcluster 4 (24.8%, 39.5%; 124 genes) and fibroblast subcluster 8 (15.9%, 49.9%; 50 genes) are *Ghr*-biased. Across all 26 subclusters the two receptors are uncorrelated (Spearman ρ = 0.23, P = 0.25; sham pseudobulk CPM). **(K)** Log2 fold change (HPS vs. Sham) of selected genes, grouped by function, in the four subclusters shown in (I). Color = unshrunken log2 fold change, clipped at ±2. Outlined cells = padj < 0.05; faded cells = tested, not significant; grey = not tested.

Interestingly, in epithelial cells, *Bcl6* (B-cell Lymphoma 6) increased (log2FC = +3.26, padj = 7.1e-30) and *Cish* (Cytokine-Inducible SH2-containing protein) decreased (log2FC = −2.91, padj = 3.1e-14) in HPS blastemas (**Fig**. 2E). In other systems, BCL6-mediated repression can block cell differentiation and promote cell proliferation, and PRL/GH are known to act via STAT5 to suppress *Bcl6* expression (*19–21*). Elevated *Bcl6* expression in this setting is therefore consistent with a loss of pituitary-derived STAT5-mediated repression. On the other hand, *Cish* expression can be induced by PRL/GH/STAT5 signaling and therefore may serve as a readout for pituitary hormone signaling in the limb(*22–24*). In the fibroblast population, *Cish* is again decreased in the HPS condition (log2FC −0.71, padj 2.1e-4) (**Fig**. 2F). Canonical hormone receptor transcript abundance did not change across cell types and pituitary hormone transcripts were largely absent from the blastema (**Fig.** S5A,B). GSEA of the HPS-responsive transcriptional program showed that fibroblasts upregulated cell-cycle/DNA-replication programs while downregulating genes involved with oxidative metabolism and protein-folding/secretory pathways. Epithelial cells showed upregulation of lymphocyte-activation programs and downregulation of RNA-processing pathways (**Fig.** S6A). These data show that HPS altered the transcriptional programs of epithelial cells and fibroblasts in distinct ways, and that pituitary input coordinates aspects of the local regenerative program during limb regeneration.

We then asked whether the relevant signaling changes might coincide with the level of receptor expression across cell types. Pituitary hormone receptors were variably distributed across cell types within the limb (**Fig.** S7). Interestingly, *Prlr* and *Ghr* expression were enriched primarily in epithelial cells and fibroblasts (**Fig.** 2G). We therefore decided to perform further subclustering of these two cell types to increase the resolution of cell type subpopulations. We reclustered epithelial cells into 16 subclusters (**Fig.** S8A). Fibroblasts were reclustered into 15 subclusters (**Fig.** S8B). We then performed differential gene expression analysis between conditions within each subcluster. Two subclusters in both epithelial cells and fibroblasts accounted for the majority of DEGs: epithelial subcluster 5 (sc5) (217 genes), epithelial sc4 (124 genes), fibroblast sc2 (178 genes) and fibroblast sc8 (50 genes) (**Fig**. 2H, H’). These four populations were transcriptionally distinct (**Fig.** S8C, D). Epithelial sc5 is a population of small secretory cells (*Fcgbp*, *Otog-like*, *Muc4*). Epithelial sc4 is likely a population of proliferating suprabasal epithelium (*Mki67*, *Gmnn, Cldn1*). Fibroblast sc2 is a subpopulation of cycling fibroblast-like blastema cells (*Prrx1*+, *Kazald2(high), Mxra5, Mki67(high)*). Fibroblast sc8 resembles a previously identified putative fibro-adipogenic progenitor (FAP) limb cell population (*Pdgfra+, Osr1+*)(*25*). To provide a possible mechanistic link between hypophysectomy and the DEG disparity between subclusters, we investigated *Prlr* and *Ghr* expression across these four populations. Interestingly, the two subclusters with the most DEGs were both *Prlr*-biased. We found that in epithelial sc5, *Prlr* and *Ghr* transcripts were expressed in 44% and 26% of cells, respectively (**Fig**. 2I). Epithelial sc4 expressed *Prlr* and *Ghr* transcripts in 25% and 39% of cells, respectively. Among fibroblasts, the separation was wider, with sc2 expressing *Prlr* and *Ghr* in 27% and 5% of sham cells and sc8 expressing *Prlr* and *Ghr* in 16% and 50% of sham cells (**Fig**. 2I). Plotting the percentage of *Prlr*- and *Ghr*-expressing cell clusters, we show that epithelial sc5 and fibroblast sc2 are *Prlr*-biased, while epithelial sc4 and fibroblast sc8 are *Ghr*-biased (**Fig.** 2J).

The epidermal populations showed the strongest STAT5 response as a result of HPS. In the proliferating suprabasal layer (epithelial sc4), *Cish* fell sharply (log2FC = −3.33, padj = 2.2e-16), accompanied by reduced ribosome biogenesis (*Nolc1*, log2FC = −0.74, padj = 0.001; *Fbl*, log2FC = −0.64, padj = 2.9e-5) and ER chaperone expression (*Hspa5*, log2FC = −1.09, padj = 0.003) (**Fig**. 2K). The small secretory cells (epithelial sc5) showed the largest change in STAT5 targets: *Cish* decreased (log2FC = −4.89, padj = 3.3e-12) and *Bcl6* increased (log2FC = +4.38, padj = 8.6e-23). Glucocorticoid- and steroid-responsive genes also fell, including *Hsd11b2* (log2FC = −2.37, padj = 4.6e-9), *Srd5a2* (log2FC = −2.06, padj = 1.2e-4). *Sesn1* also decreased (log2FC = −2.31, padj = 1.5e-8). Both epithelial sc4 and sc5 increased *Igf1r* (sc4: log2FC = +0.57, padj = 0.039; sc5: log2FC = +0.87, padj = 0.016), possibly consistent with compensation for reduced ligand. In epithelial sc5, we observed increases in *Muc5b* (log2FC = +3.72, padj = 1.1e-11) and innate defense gene *Nos2* (log2FC = +1.90, padj = 0.003), along with reduced *Fgfbp1* (log2FC = −1.16, padj = 4.6e-9).

Fibroblast sc8 showed a distinct decrease in hormone signaling readouts. HPS reduced *Cish* (log2FC = −1.73, padj = 7.8e-4) and the GH-induced binding protein *Igfbp3* (log2FC = −1.82, padj = 8.4e-8) (**Fig**. 2K). The thyroid hormone receptor *Thra* decreased (log2FC = −1.66, padj = 8.7e-4) while *Dio3* increased (log2FC = 1.34, padj = 0.027), consistent with altered thyroid signaling after loss of TSH. Beyond hormone signaling, sc8 downregulated the mature matrix genes *Fbn1* (log2FC = −1.34, padj = 0.007) and *Col4a1* (log2FC = −1.31, padj = 0.046). *Fgf7*, which encodes a paracrine mitogen for epithelium, decreased as well (log2FC = −1.52, padj = 0.013). At the same time, sc8 increased expression of *Runx1* (log2FC = +2.00, padj = 8.4e-8), which has been shown to be linked to activation of fibroblasts after injury and fibrosis. Without pituitary input, these FAPs therefore lose a local GH–IGF program while possibly adopting a more fibrotic state. Fibroblast sc2 responded differently. Its response was largely directional, with 147 of 178 DEGs decreasing. These cells decreased collagen and secretory machinery, including *Col1a1* (log2FC = −0.46, padj = 0.009), *Serpinh1* (log2FC = −0.74, padj = 5.3e-4), *Lox* (log2FC = −0.39, padj = 0.012), and *Sparc* (log2FC = −0.37, padj = 0.012). Cytoskeletal genes decreased such as *Tagln* (log2FC = −0.89, padj = 5.3e-4), *Lgals1* (log2FC = −0.57, padj = 5.3e-7) and *Crip1* (log2FC = −0.69, padj = 0.001).

These findings point to cell type-specific responses to pituitary loss, concentrated in cells equipped to receive pituitary hormone signals rather than a generic tissue-wide effect. The epidermis shifted from a hormone-responsive barrier toward a mucous-producing or defensive state. FAP-like cells lost a local GH–IGF program. *Kazald2-*expressing blastema progenitors maintained proliferation markers but decreased their expression of matrix and cytoskeletal genes.

### Local PRLR signaling is required for blastema participation and digit morphogenesis and can be restored by local hormone delivery

*Prrx1*+/*Kazald2*(high) fibroblast cells, known to be important during blastema formation (*26*), highlighted a bias toward *Prlr* over *Ghr*. Therefore, we wanted to determine whether PRLR acts locally within the regenerating limb. *Prlr* expression was locally enriched in the wound epidermis at 1 dpa (**Fig**. 3A). We detected PRL1 protein directly in regenerating limb tissue across early regeneration (**Fig**. 3B). To directly test whether *Prlr* is required locally within limb cells during regeneration, we used a translation-blocking morpholino knockdown strategy to reduce *Prlr* levels. We knocked down *Prlr* before amputation and then re-administered the morpholino or morpholino control at 7 dpa. Morphology and blastema size were evaluated throughout the regeneration time course (**Fig**. 3C). By 31 dpa, *Prlr* knockdown limbs showed a significant reduction in the number of distinct digit outlines compared to controls (p = 0.0001) (**Fig**. 3D, E).

**Figure 3.**
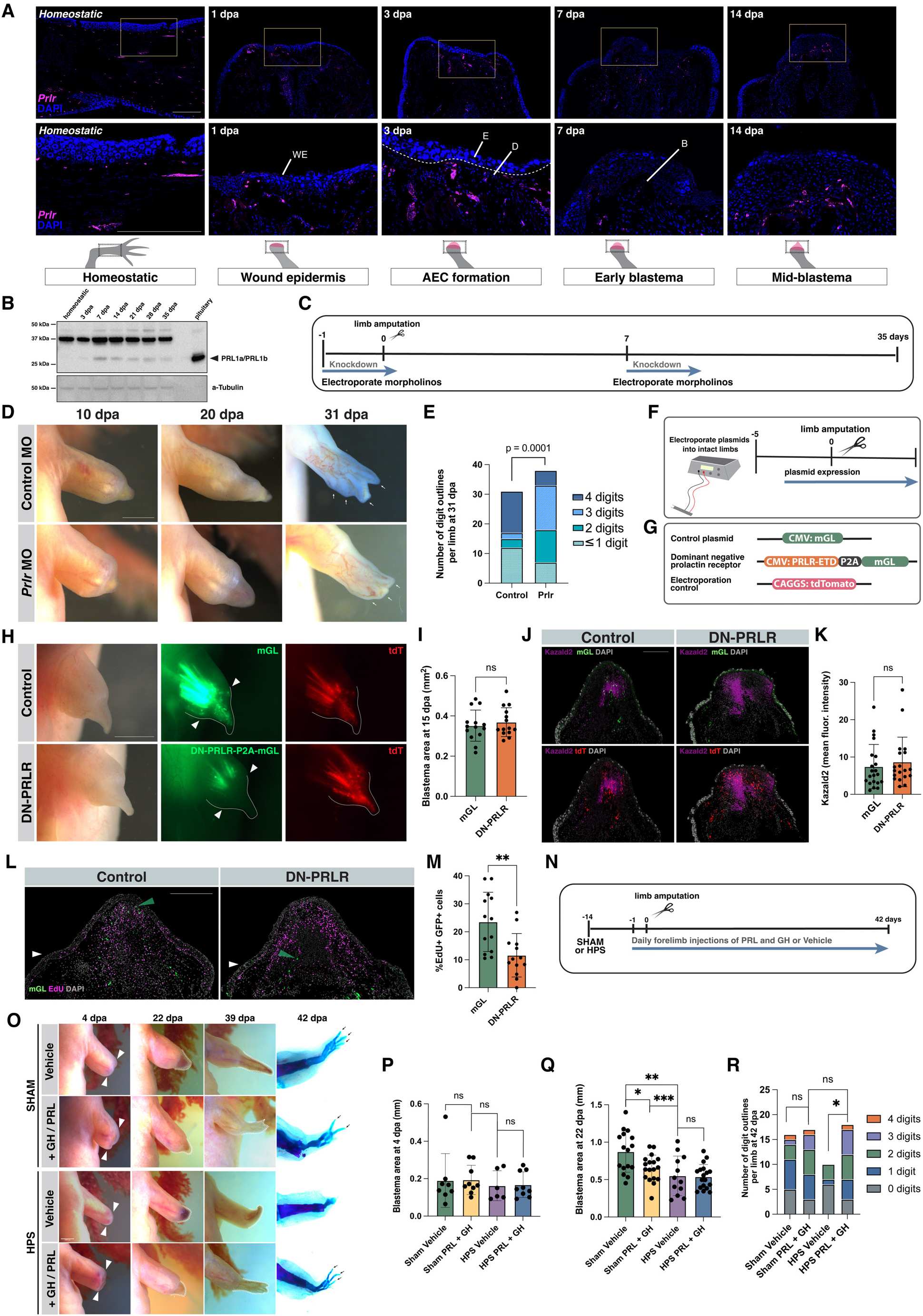
Local PRLR signaling is necessary for digit morphogenesis and local hormone replacement rescues regeneration in hypophysectomized limbs. **(A)** *Prlr* HCR-FISH time course through limb regeneration, showing where *Prlr* transcripts localize, with DAPI in blue, at Homeostatic, 1, 3, 7, and 14 dpa. The yellow box is depicted as a higher magnification below. The regeneration stage (Homeostatic, Wound epidermis, Apical epidermal cap formed, Blastema formation, Mid-blastema) and the location of section are represented below each timepoint. WE = wound epidermis, E = epidermis, D = dermis, B = blastema. Scale bar, 500 µm. (B) Western blot of total protein from limbs at Homeostatic, 3, 7, 14, 21, 28, and 35 dpa, with a blank lane and pituitary as a positive control, probed for PRL1A/B and alpha tubulin as a loading control. (C) Morpholino electroporation timeline and diagram: morpholinos were electroporated 1 day before and 7 days after amputation (knockdown), followed to 35 days. (D) Control and *Prlr* morpholino limbs at 10, 20, and 31 dpa. Arrowheads mark digit outlines. Scale bar, 2 mm. Images were acquired ventrally through 24 dpa and dorsally at 31 dpa.(E) Digit number at 31 dpa was reduced by *Prlr* morpholino treatment (control, n = 34 limbs; *Prlr* MO, n = 38 limbs). Fisher–Freeman–Halton exact test, p = 0.0001 **(F)** Electroporation and injection timeline for dominant-negative prolactin receptor and control plasmids. **(G)** Control, truncated construct of prolactin receptor including extracellular and transmembrane domains but lacking the intracellular signaling domain (dominant-negative prolactin receptor, DN-PRLR), and tdTomato electroporation-control plasmid design. **(H)** Brightfield, mGL, and tdTomato images of control and DN-PRLR blastemas at 15 days post-amputation (dpa). Scale bar, 1 mm. **(I)** Blastema area at 15 dpa (n = 14 animals per group; not significant). Unpaired two-tailed Welch’s t-test. **(J)** Sections showing mGL, tdTomato, and the blastema marker *Kazald2* in control and DN-PRLR blastemas. Scale bar, 500 µm. (K) *Kazald2* mean fluorescence intensity quantification. Unpaired two-tailed Welch’s t-test. **(L)** EdU labeling showing loss of DN-PRLR-expressing cells from the blastema without reduced proliferation. Scale bar, 500 µm. **(M)** Percentage of EdU-positive nuclei among mGL- and DN-PRLR-electroporated cells (n = 26 limbs for control, n = 25 limbs for DN-PRLR). Unpaired two-tailed Welch’s t-test, p= 0.0035. **(N)** PRL1 and GH rescue experimental design. Animals underwent HPS/sham surgery and were allowed to recover for 14 days. Daily double forelimb injections with GH and PRL1 or vehicle (saline) injection for 42 days post amputation. **(O)** Representative Sham and HPS vehicle and GH and PRL1-treated HPS and Sham limbs in brightfield at 4, 22, and 39 dpa. Amputation plane marked with white arrowheads. Regenerated region outlined in white dashed line. Alcian blue/Alizarin red skeletal stain at 42 dpa for each limb. Digits formed marked with black arrowheads. Scale bar, 1000µm. **(P)** Blastema area at 4 dpa (n= 18 limbs GH and PRL HPS and sham-operated, n=16 limbs vehicle sham-operated, n= 10 limbs vehicle HPS-operated; Unpaired two-tailed Welch’s t-test, not significant). **(Q)** Blastema area at 22 dpa (right) (n= 18 limbs GH and PRL HPS and sham-operated, n=16 limbs vehicle sham-operated, n= 10 limbs vehicle HPS-operated; Unpaired two-tailed Welch’s t-test, p = 0.0107, 0.0044, 0.0003). **(R)** Number of digit outlines per limb at 42 dpa, Vehicle HPS-treated had significantly less digits at 42 dpa compared to GH and PRL HPS-treated limbs (n= 18 limbs GH and PRL HPS and sham-operated, n=16 limbs vehicle sham-operated, n= 10 limbs vehicle HPS-operated; Fisher–Freeman–Halton exact test, p= 0.0418).

To test how disrupting *Prlr* signaling affects individual cells’ contributions to the blastema, we electroporated intact limbs with either a dominant-negative PRLR construct with a fluorescent reporter (DN-PRLR-mGL) or a control plasmid expressing mGL alone. All limbs were also co-electroporated with a tdTomato electroporation control (**Fig**. 3F-G). After confirming comparable mGL and tdTomato expression, limbs were amputated and imaged throughout regeneration. In the control, tdT cells (red) and mGL+ cells (green) spread well into the distal blastema, past the amputation plane. However, in the DN-PRLR-mGL group, tdT (red) still appeared in the blastema at levels similar to the control, but electroporated cells (green) were absent from the distal blastema. DN-PRLR-mGL electroporated cells largely failed to pass the amputation plane into the blastema (**Fig**. 3H). At 15 dpa, blastema area did not differ between DN-PRLR-mGL and control limbs (**Fig**. 3I). Moreover, expression of the blastema marker *Kazald2* was not affected within the blastema itself (**Fig**. 3J, K). Local disruption of *Prlr* signaling therefore did not prevent blastema formation or early growth. In contrast, proliferation in blastemas was significantly reduced in DN-PRLR-mGL positive cells (as indicated by %EdU-positive nuclei) compared with mGL-positive control cells (**Fig**. 3L, M). Together with the morpholino knockdown results, these data suggest that PRLR signaling is required cell-autonomously at the earliest stages of blastema formation and for digit morphogenesis via signaling within the limb itself.

This led us to evaluate whether directly restoring ligand availability at the wound site in HPS animals would be sufficient to rescue regeneration. Since both human GH and PRL are known to be capable of signaling through PRLR (*27*), we administered human GH and axolotl PRL1a/PRL1b or vehicle by daily intralimb injection for 42 days directly below the amputation site in HPS or sham-treated animals (**Fig**. 3N, O). The blastema area was measured until digits began to form. As expected, the blastema area between HPS-treated and sham-treated animals, both injected with vehicle, was significantly different at 22 dpa (p= 0.0044) (**Fig.** 3P-R). At 42 dpa, limbs were harvested from animals, and skeletal patterning was analyzed via Alcian blue and Alizarin red staining. Animals that underwent HPS and were injected with GH and PRL1 had more digits formed by 42 dpa compared to the vehicle controls (p=0.0418). Considered together, these three perturbations provide evidence that PRLR/GHR signaling at the wound site is necessary for normal axolotl limb regeneration.

### Pituitary dependence is conserved in fish fin and mammalian digit tip regeneration

Finally, we asked whether this pituitary/PRLR-dependent axis is specific to axolotl limb regeneration or represents a more broadly conserved vertebrate mechanism. In tilapia, caudal fin amputation (**Fig**. 4A-C) drove position-specific changes in the same signaling components identified in the axolotl blastema. Specifically, *Prlr2, Stat5, Stat1a,* and *Cish* were each altered in proximal versus distal fin tissue 24 hours after amputation (**Fig**. 4D-F, S9A-E). This indicates that amputation triggers a local, spatially organized JAK-STAT and PRLR transcriptional response in an independent vertebrate lineage that diverged over 400 million years ago [21]. Consistent with the axolotl phenotype, HPS tilapia showed significantly reduced caudal fin regeneration at 17 dpa compared with sham controls (**Fig**. 4G-I), demonstrating that pituitary function is required for complex structural regeneration in a second, distantly related vertebrate.

**Figure 4.**
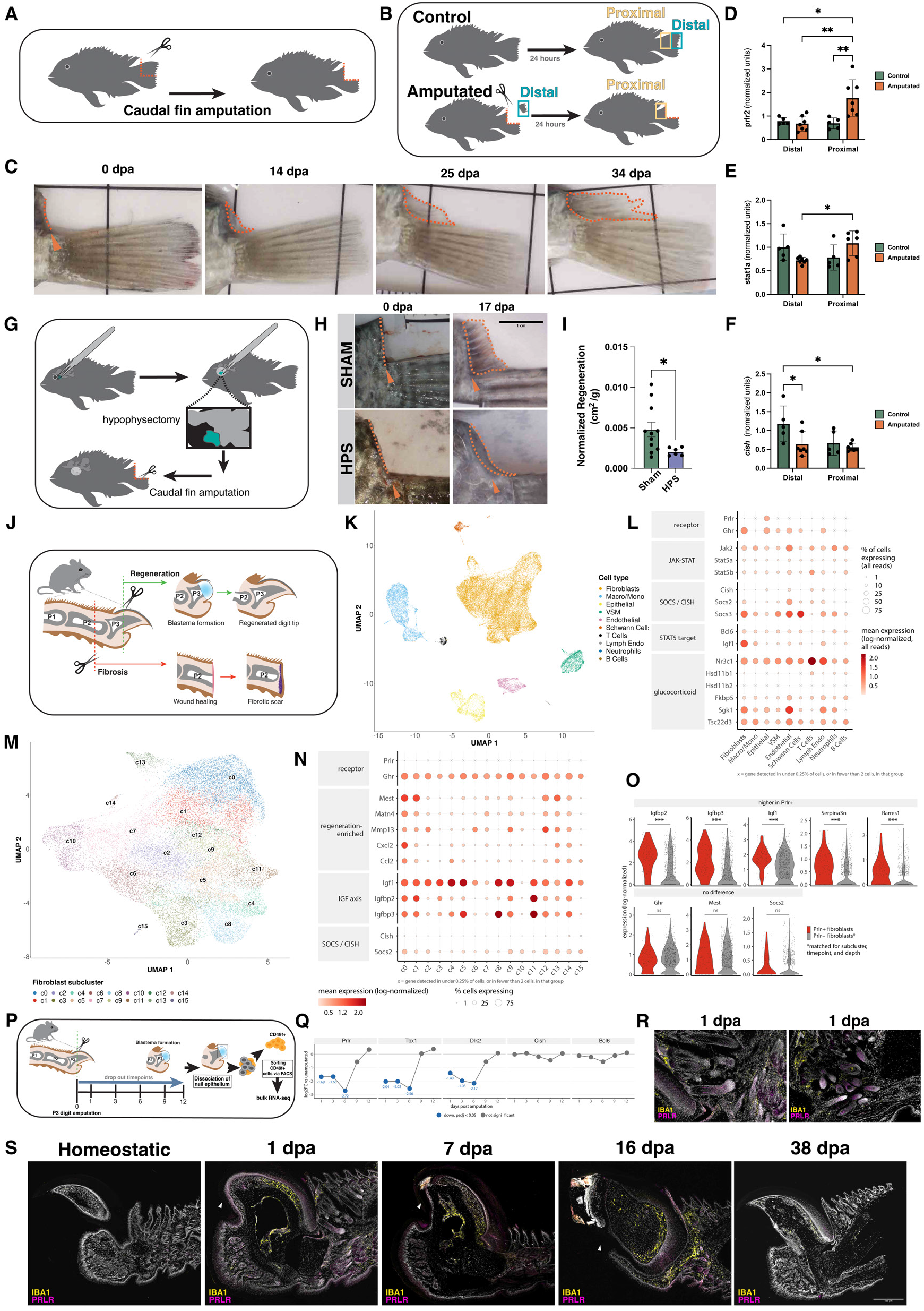
Evolutionarily conserved and comparative insights into appendage regeneration. **(A)** Schematic of the caudal-fin amputation procedure in tilapia. **(B)** Sampling design for qPCR: Control and Amputated fish were each sampled at Distal and Proximal fin-clip positions post-procedure. **(C)** Representative brightfield images of the caudal-fin regeneration time course after amputation (0, 14, 25, 34 dpa); dashed orange line marks the regenerate outline. **(D)** P*rlr2* (qPCR). Values are normalized to ef1a and depicted as arbitrary units. Bars show mean ± SD. Significance was assessed using two-way ANOVA and Tukey’s multiple comparisons test. *: p < 0.05, ** p < 0.01. Comparisons not shown were not significant. **(E)** *Stat1a* (qPCR). Values are normalized to ef1a and depicted as arbitrary units. Bars show mean ± SD. Significance was assessed using two-way ANOVA and Tukey’s multiple comparisons test. *: p < 0.05. Comparisons not shown were not significant. **(F)** *Cish* (qPCR) Values are normalized to ef1a and depicted as arbitrary units. Bars show mean ± SD. Significance was assessed using two-way ANOVA and Tukey’s multiple comparisons test. *: p < 0.05. Comparisons not shown were not significant. **(G)** Schematic of hypophysectomy in tilapia (pituitary removed via the eye) followed by caudal-fin amputation. **(H)** Representative images of Sham vs. hypophysectomized (HPS) fish fins at 0 and 17 dpa; orange arrowheads mark the amputation plane. Scale bar = 1 cm. **(I)** Quantification of normalized regenerated area (cm^2^/g) at 17 dpa, Sham vs. HPS. Individual animals plotted with bar = mean ± error bar; unpaired t-test, *: p < 0.05. **(J)** Schematic of the mouse digit-tip amputation model: P3 amputation with blastema formation and regenerated digit tip; P2 amputation with failed regeneration and fibrosis. Adapted from Jou et al., 2024. **(K)** UMAP of 43,408 cells from unamputated (UA), 11, 12, 14, 17, and 28 dpa P3 mouse digit tips (reanalyzed from scRNA-seq data from [**28**, **29**, **30**]), colored by published cell type. **(L)** Dot plot of prolactin and growth hormone pathway genes by cell type. Color represents mean log-normalized expression; x = detected in <0.25% of cells or in fewer than 2 cells; dot size represents percentage of cells detecting the gene. **(M)** UMAP of 29,393 P3 fibroblasts across 16 subclusters. **(N)** Dot plot of receptor, regeneration-enriched fibroblast marker (*29*), IGF axis, and SOCS/CISH genes by fibroblast subcluster. Formatted equivalent to (L). **(O)** Expression of selected genes in *Prlr*+ fibroblasts (86 cells) and *Prlr*-fibroblasts (847 cells) matched for subcluster, timepoint, and sequencing depth. 21 genes were identified as differentially expressed (19 higher in *Prlr+* cells; Wilcoxon test; detected in ≥25% of cells; log2 fold change > 0.58; *** = padj <0.001; ns = not significant). **(P)** Schematic of bulk RNA-seq of FACS-sorted *CD49f*+ cells from P3 nail epithelium at UA, 1, 3, 6, 9, and 12 dpa (n = 3 per timepoint). **(Q)** Expression of selected genes in *CD49f*+ cells, shown as log2 fold change versus UA (DESeq2 Wald test). Blue = decreased (padj < 0.05); gray = not significant. (**R**) High-magnification immunofluorescence of PRLR (magenta) and IBA1 (yellow) around hair follicles, showing PRLR signal localized to follicle structures. **(S)** Immunofluorescence of the mouse P3 digit regeneration time course (1, 7, 16, 38 dpa) for IBA1 (yellow) and PRLR (magenta); white arrowheads indicate the amputation plane. Scale bar 500µm.

We then asked whether elements of this pathway were detectable and applicable in the mouse digit tip model of mammalian regeneration. Amputation through the P3 digit region leads to spontaneous regeneration, whereas amputation through the P2 region leads to fibrosis (**Fig**. 4J). We first confirmed that our P3 model followed the expected regenerative trajectory by tracking morphology and Masson’s trichrome histology (**Fig.** S10A, B). This established the tissue- and skeletal-remodeling timeline against which the endocrine transcriptional data were interpreted. Re-analysis of a published scRNA-seq profile of P3 digits across the regenerative time course generated the expected cell types (**Fig.** S11). *Prlr* was detected in 41.7% of epithelial cells and 0.4% of fibroblasts and *Ghr* was detected in 52% of epithelial cells and 63% of fibroblasts. (**Fig**. 4K, L) (*28–30*). Reclustering the fibroblasts revealed a subcluster, c1 (*Mest+, Igfbp2-high)* that was largest at 11-14 dpa and contained 34 of 107 *Prlr*+ fibroblasts (1.07% of c1 cells compared to 0.36% of all fibroblasts)(*29*). At 14 dpa, the *Prlr+* enrichment was 3.8 times more than in other fibroblast clusters (**Fig**. 4M,N). Compared with *Prlr-*fibroblasts, *Prlr*+ cells were higher in *Igf1*, *Igfbp2* and *Igfbp3* (**Fig.** 4O). Although low in number, this suggests that Prlr marks an IGF-binding-protein-high state within the regenerating tissue. Moreover, these data suggest that pituitary signaling can act directly on the cell population that drives regrowth. To investigate the epithelial population, we analyzed bulk RNA-seq of CD49f+ nail epithelium through early regeneration (**Fig.** 4P). *Prlr* decreased at 1-6 dpa and returned to baseline at 9-12 dpa (**Fig.** 4Q, S12). To confirm that PRLR protein is present in the regenerating digit tissue, we supplemented the transcript-level receptor data from scRNA-seq (**Fig.** 4L, N, Q), by performing immunofluorescence staining for IBA1 (macrophage marker) and PRLR (**Fig**. 4R, S). PRLR localization was concentrated at the hair follicles in homeostatic digits, which is consistent with previous studies in mammals (*31*). IBA1 served as a positive control, showing upregulation at the injury site and during regeneration before returning to baseline levels. We validated that the scRNA-seq data accurately predicted the observed robust PRLR in epithelial cells at 1 and 7 dpa, and within proximal connective tissues at 16 dpa. Together, these data establish that the receptor is present and positioned locally to respond to pituitary signaling in the digit.

## Discussion

Our findings connect pituitary function to the cellular responses that support appendage regeneration across multiple vertebrate lineages. Surgical pituitary removal impaired blastema growth in axolotl and fin regeneration in tilapia, while local disruption of PRLR signaling compromised blastema cell participation and digit formation in axolotl. Together with the pituitary’s sustained transcriptional response to limb loss, these findings identify an endocrine contribution to regeneration that links a distant organ with local tissue repair. The shared requirement for pituitary function in axolotl limb and tilapia fin regeneration suggests that endocrine support of appendage regeneration may have deep roots, across 400 million years of vertebrate evolution.

The pituitary’s transcriptional response to limb loss highlights its role as an injury-responsive regulator of distant regeneration. Through an unknown mechanism following limb amputation, the axolotl pituitary engages in transcriptional programs associated with cell activation, developmental signaling, and the upregulation of genes typically associated with adult pituitary stem cells. Multiple studies in mammals have shown that SOX2+ pituitary stem cells are capable of regenerating the gland’s tissue after direct injury or targeted cell ablation (*32–35*).

Furthermore, regulation of this stem cell niche is controlled by developmental pathways including FGFs and Wnt (*36, 37*), which we observed to increase after limb amputation. It has not been previously shown that adult pituitary stem cells can respond to distant injury of non-endocrine tissue. In highly regenerative organisms, these signaling pathways may be activated in response to circulating inflammatory markers, acute pituitary damage, or to compensate for increased demand for pituitary hormone output. Further investigation into this transcriptional response to distant injury is an exciting avenue for future experimentation.

The pronounced epithelial and fibroblast response to pituitary removal suggests that endocrine input may be particularly important for wound epidermis function, control of fibrosis, blastema participation, and/or cellular differentiation and digit patterning. Increased *Bcl6* and decreased *Cish* expression were consistent with reduced STAT5-associated signaling, while the concentration of transcriptional changes in subpopulations with distinct *Prlr* expression suggests that cellular state and receptor availability may shape responses to pituitary input. This provides a potential mechanism through which circulating hormones could coordinate distinct functions within regenerating tissue. Differences in receptor expression and cellular state could enable distinct populations to interpret a shared hormonal environment differently, linking systemic endocrine input to the diverse cellular activities required for regeneration. For example, pituitary signals might support wound epidermis function in epithelial cells while influencing matrix remodeling and blastema participation in fibroblasts. Such a model would allow circulating hormones to coordinate complementary responses across cell populations without requiring a uniform effect throughout the regenerate. Although hypophysectomy may also influence regeneration through systemic physiological changes, the local PRLR perturbations independently support a requirement for hormone responsiveness within limb cells. Determining how epithelial responses contribute to blastema growth will require cell-type-specific manipulation of these pathways.

Our finding that DN-PRLR-expressing cells fail to enter the blastema indicates a cell-autonomous requirement for PRLR signaling in blastema cell specification or migration, or both. Notably, *Tagln* expression was down in HPS in fibroblast sc2 – the most *Prlr*-biased and most transcriptionally responsive fibroblast population (**Fig.** 2K). In an existing amphibian limb regeneration dataset, *Tagln* is a marker of mature fibroblasts, and these fibroblasts undergo transcriptional changes during blastema formation consistent with dedifferentiation (*26*). Thus, pituitary loss may result in stump cells becoming “stuck” in a differentiation state that is ordinarily transient during regeneration. Alternatively, in the absence of pituitary signals, fibroblasts may have dedifferentiated or otherwise activated normally in response to amputation, but then failed to rebuild cytoskeletal machinery needed for a later migratory step, a process which shares similarity to epithelial-to-mesenchymal transition (EMT) (*38*). In human and mouse tumors, Transgelin appears to play such a role, serving as a canonical marker of myofibroblastic cancer-associated fibroblasts (myCAFs) (*39, 40*). Because myCAFs are associated with the invasive front of tumors during metastasis, reduced *Tagln* in regenerating axolotl limbs in the HPS context could indicate that fibroblast-derived progenitor cells cannot properly migrate into the blastema. A shared feature between cancer and regeneration may be the acquisition of fibroblast states that support tissue remodeling and cell movement, which regeneration leverages to rebuild tissue and tumors use to facilitate invasion. Whether PRLR signaling supports such a state during regeneration remains unresolved, as the exclusion of DN-PRLR-expressing cells could also reflect altered survival, differentiation, or retention. Tracking these cells would help determine whether endocrine input enables blastema participation by regulating their movement or state.

Combined GH and PRL1 administration partially restored regenerative capacity in HPS axolotls, increasing digit formation despite loss of the gland. This conclusion is further supported by gene sets related to generalized nutrition or illness and to starvation and autophagy. These gene sets were predominantly decreased rather than increased in HPS tissue (Fig. S6B), the opposite of what would be expected if HPS animals were malnourished. Although broader physiological effects of HPS cannot be excluded, these transcriptional findings, together with local hormone replacement and PRLR perturbation, support a functional role for pituitary hormone signaling in the regenerating limb. The relative contributions of GH and PRL, and whether their combined effects reflect complementary or overlapping functions, remain to be determined.

Rather than a salamander-specific innovation, pituitary-dependent regulation of regeneration appears to be an ancestral vertebrate feature, retained since before the divergence of ray-finned and lobe-finned lineages. The coordinated trajectory of *Prlr, Igfbp2,* and *Igfbp3* in mouse P3 fibroblasts after amputation suggests that pituitary hormones can act directly on regeneration-driving cell populations. These data indicate that components of the same endocrine signaling axis identified in axolotl and tilapia are differentially engaged between regenerative and fibrotic outcomes in a mammalian system. Our data raise the possibility that enhancing endocrine signaling, for instance by increasing pituitary hormone receptors at the wound site or local pituitary hormone concentrations, could offer an independent and complementary route to unlocking regenerative competence in mammalian tissue.

## Materials and Methods

### Animals

All axolotl experimentation was approved and conducted in accordance with Harvard University’s Institutional Animal Care and Use Committee (IACUC). All mouse experimentation was approved and conducted in accordance with Harvard University’s IACUC or Brigham and Women’s Hospital IACUC. All tilapia experimentation was approved and conducted in accordance with University of Hawai’i IACUC.

Wild type leucistic axolotls were housed individually in 2.63 g/L Instant Ocean Water with 4000µS conductivity at 18°C. Animals were fed axolotl sinking soft moist pellets. In all experiments, clutch-mate animals were randomly assigned to each experimental group. Axolotls used in experiments were 6 months-1 year old, 8-15 cm tail-to-snout length.

Mozambique tilapia (*Oreochromis mossambicus*) of both sexes were reared at the Tuahine Aquaculture Research and Education Center, University of Hawaiʻi at Mānoa in fresh water (FW; 0.1 ± 0.1‰, municipal water) as previously described (*41*). Ahead of trials, fish were kept in outdoor 700-L tanks filled to 375-L under natural photoperiod and fed 1.5 mm Aquaxcel (Star Milling, Perris, CA) twice daily *ad libitum*.

For mouse digit histology experiments, C57BL/6 mice (strain code 027) were obtained from Charles River Laboratories. The animals were maintained in the Harvard Biological Research Infrastructure facility. Mice were housed in groups of up to five animals per cage. Each cage contained shredded paper, red tubes, and access to food and water. Animals were randomly divided into treatment groups. Equal numbers of male and female mice (aged 5-8 weeks) were used for all experiments. For mouse nail epithelium RNA bulk sequencing experiments, 6-week-old female FVB/NJ mice (JAX #1800) were used. The animals were housed in the Hale vivarium maintained by the Brigham and Women’s Hospital (BWH) Center for Comparative Medicine.

### Experimental Design

#### Surgical techniques

Hypophysectomy, amputation, and injections

##### Axolotl

Removal of the pituitary gland (hypophysectomy) was carried out on axolotls sized 8-15 cm tail-to-snout length. Animals were anesthetized with 0.1% (w/v) tricaine before any procedures. After confirming anesthesia, the axolotl was placed dorsally and a small incision was made in the lateral oral cavity. A small cartilage flap on the roof of the mouth was removed to access bone. Utilizing a dissecting microscope, Dumont forceps and fine scissors were used to cut a window into the roof of the mouth in the area surrounding the pituitary. The pituitary was removed using forceps and light suction. Sham operations received the same treatment except for pituitary removal. Axolotls recovered overnight in 0.5% (w/v) sulfamerazine. All limb amputations were performed at the mid-stylopod and the bone was then trimmed back from the amputation plane. Growth hormone (GH: stock 0.1mg/mL) and prolactin (PRL; stock 2.5mg/mL) were administered by daily intralimb injection into both forelimbs beginning 1 day prior to amputation and 13 days post sham or hypophysectomy surgery. Dosing was targeted to 1.2µg GH and 2.5µg of PRL per limb per day (both forelimbs amputated and injected). Digit outcomes were scored at the same endpoint.

##### Tilapia

First, 12 adult tilapia 17.7 ± 1.9 g of mixed sexes were tagged with passive integrated transponder (PIT) tags (HPT8, Biomark, Boise, ID) and transferred to 45 L FW (0.1 ± 0.1‰) aquaria for 5 days prior to caudal fin amputation. Feeding was withheld 24 h prior to caudal-fin amputation. On the day of amputation, seven fish were anesthetized with 2-phenoxyethanol (2-PE; 0.6 mL/L, Sigma-Aldrich, St. Louis, MO) and subjected to caudal fin amputation, while the other 5 fish were kept intact as controls. Immediately following amputation, the salinity was increased to brackish water (BW; 12 ± 0.1‰) with sea salt (Instant Ocean, Blacksburg, VA). At the time of amputation, the distal fin clip was collected from the amputated group. At 24 h, proximal caudal fin tissue was collected from the amputated group, while distal and proximal tissue were collected from the intact control group (Fig. 4B). All tissue were immediately preserved in RNA*later* (Invitrogen, Carlsbad, CA) for qPCR analysis. To monitor fin regeneration rates, an additional 9 PIT-tagged fish (5 amputated and 4 control fish) were photographed with a digital microscope (TM4K-AF Max, TOMLOV) at 0, 14, 25 and 34 days post amputation (Fig. 4C). In a follow-up trial, 25 PIT-tagged tilapia of mixed sexes weighing 68.6 ± 4.0 g were divided into 4 tanks (6-7 fish per tank), supplied with 375 L of FW. Water in each tank was recirculated with independent canister filters (Fluval FX-2) and kept at 23 - 27 °C. Two weeks following tagging, the salinity of all tanks was adjusted to BW followed by hypophysectomy (n=12) or sham operation (n=13) the following day, then caudal fin amputation four days after hypophysectomy. Feed was withheld 24 h prior to hypophysectomy and throughout the 17 days. Fish were photographed after caudal fin amputation on day 0 and 17 days post amputation. Regeneration was measured on day 17 using ImageJ (*42*) and normalized by calculating the ratio of regenerated area (cm^2^) to body weight (g)

Removal of the pituitary (hypophysectomy) was carried out in male and female tilapia weighing 68.6 ± 4.0 g and measuring 12-15cm in total length. Procedure as cited in (*43, 44*). Caudal fin amputations were performed within 1 cm of the endochondral plate with the upper half of the caudal fin removed as modified from (*45*). After surgery, all fish were returned to their original tanks and exposed to the antibiotic Kanaplex (0.2% kanamycin sulfate) for 5 days.

##### Mouse

For histology experiments, mouse P3 digit amputation was completed as described in (*28*) with one modification. Only hindlimb digits 2 and 4 were amputated through the third phalangeal segment. For nail epithelium RNA bulk-seq experiments, hindlimb digits 2-5 of 6-week-old female FVB/NJ mice were amputated through the P3 segment.

### qPCR

#### Tilapia

Total RNA was extracted from distal and proximal caudal-fin tissues using TRI Reagent following the manufacturer’s protocol (MRC, Cincinnati, OH) and reverse transcribed using a cDNA reverse transcription kit (Cat. No.: 4368813; Thermo Fisher Scientific, Waltham, MA). The levels of reference and target genes were determined using the relative quantification method(*46*), in which relative expression levels and PCR efficiencies were calculated from standard curves produced from serial dilutions of tissue cDNA using a StepOnePlus real-time qPCR system (Thermo Fisher Scientific). The qPCR reactions (15 μL) contained 7.5 µL of SYBR Green PCR Master Mix (Cat. No.: 4368708; Applied Biosystems, Warrington, UK), 0.3 µL of 10 µM forward and reverse primers, and 1 μL of cDNA, with cycling parameters as follows: 95 °C for 10 min, followed by 40 cycles at 95 °C for 15 s and 60 °C for 1 min. Specific primers for *Prlr2*, *Stat1a, Aqp3, Prlr1, Ostf1* have been previously described (*43, 47–49*), and primers for *Cish*, were newly designed. *Elongation factor 1α* (*Ef1α*) was used as the reference gene to normalize target gene levels (*43*) after confirming its stable expression across treatment groups. Data are expressed as the mean fold-change ± S.E.M from the distal control samples.

### Immunofluorescence, HCR-FISH, Masson’s Trichrome, and EdU

Cryosectioned tissue fixed in 4% PFA from axolotl and mouse were rehydrated with PBS and permeabilized with 0.3% Triton-X-100 PBS. Post-permeabilization, slides were rinsed with PBS and EdU was visualized via sulfo-cyanine 3 azide. Slides were washed 3X with PBS. Slides were then stained and labeled with any additional antibodies. Primary antibodies used: anti-IBA1 (Chicken) AvesLabs IBA1-0100, anti-PRLR (Rb) Bioss bsm-61624r, and anti-PRL1 (Rb) (custom antibody DZ41666).

Sections were blocked in 10% serum from the host species of the secondary antibody in 0.3% Triton-X-100 in PBS. Primary and secondary antibodies were diluted in blocking buffer. All slides were incubated with DAPI (Sigma D9542).

HCR-FISH v3.0 was performed according to manufacturer instructions and probes designed using Probe Generator tool (Monaghan Lab, Northeastern University, MA, USA)(*50*).

Masson’s Trichrome was used according to manufacturer instructions (Polysciences product number 25088).

For skeletal preparations, limbs were fixed overnight in 95% ethanol with gentle rocking at room temperature. Limbs were dehydrated in 100% acetone overnight. Specimens were stained in working solution of Alcian blue 8GX (Sigma A5268) and Alizarin red S (Sigma A5533) at 37°C for two weeks. Specimens were then cleared through a graded 1%KOH/glycerol series.

### Morpholino-mediated knockdown of prolactin receptor during limb regeneration

A translation-blocking morpholino targeting axolotl prolactin receptor (*Prlr*) was designed and synthesized by GeneTools using the *Prlr* transcript sequence XM_069627116.1. The *Prlr*-targeting morpholino (5′-TTACTTTTAGTTTTCGTCGCATG-3′) is complementary to a 23-nucleotide region encompassing one nucleotide upstream of the translational start site and the first 22 nucleotides of the coding sequence, including the ATG start codon. The standard negative control morpholino (5′-CCTCTTACCTCAGTTACAATTTATA-3′) targets a human β-globin intron mutation associated with β-thalassemia. Morpholinos were prepared from 1 mM stocks at a nominal working concentration of 500 μM in 0.7X PBS. Fast Green was included to aid visualization during injection.

Seven-month-old sibling wild-type juvenile axolotls naïve to amputation, 5-8 cm in snout-to-tail length, were used. Animals were anesthetized with tricaine for injection, electroporation, and amputation. Both forelimbs were injected proximally with either *Prlr*-targeting or standard control morpholino and electroporated one day before amputation. Injections were performed using filament-containing borosilicate glass needles with an outer diameter of 1 mm and an inner diameter of 0.75 mm. Each animal received approximately 10 μL of injection solution in total. Following injection, electroporation was performed with animals immersed in 1X PBS using a NepaGene Super Electroporator NEPA21 Type II. The poring pulse consisted of three pulses at 150 V, with a pulse length of 5 ms, a pulse interval of 10 ms, a decay rate of 0%, and positive polarity. The transfer pulse consisted of five pulses at 50 V, with a pulse length of 50 ms, a pulse interval of 950 ms, a decay rate of 0%, and positive polarity. The distance between electrodes was approximately 3 mm.

The following day, we assessed the fluorescein using a Leica microscope and amputated both forelimbs proximally through the electroporated region. Animals were maintained overnight in 0.5% (w/v) sulfamerazine solution and returned to fresh axolotl water at 1 day post-amputation (dpa). At 7 dpa, regenerating blastemas underwent a second round of morpholino injection and electroporation.

### Misexpression of dominant-negative prolactin receptor during limb regeneration

Wild-type juvenile axolotls naïve to amputation, 5-8 cm in snout-to-tail length, were used to examine the effects of dominant-negative prolactin receptor (PRLR) expression on limb regeneration. The dominant-negative construct, pCMV-PRLR-ETD-P2A-mGL(Tol2), encoded PRLR-ETD and an mGL fluorescent reporter separated by a P2A sequence. The pCMV-mGL(Tol2) plasmid was used as the control. Plasmid injection solutions were prepared at a nominal concentration of 400 ng/μL in 0.7X PBS, with Fast Green added to aid visualization during injection.

Both hindlimbs were electroporated with either the dominant-negative or control plasmid, together with pCAG-tdTomato as an additional electroporation reporter, with each plasmid prepared at a concentration of 400 ng/μL. Following injection, animals were immersed in cold 1X PBS, and electroporation was performed using a NepaGene Super Electroporator NEPA21 Type II. The poring pulse consisted of three pulses at 150 V, with a pulse length of 5 ms, a pulse interval of 10 ms, a decay rate of 0%, and positive polarity. The transfer pulse consisted of five pulses at 50 V, with a pulse length of 50 ms, a pulse interval of 950 ms, a decay rate of 0%, and positive polarity. The distance between electrodes was 3 mm.

Following assessment of fluorescent reporter expression, both hindlimbs were amputated proximally through the fluorescent region. Animals were maintained overnight in sulfamerazine solution and returned to fresh axolotl water the following day. Regenerating limbs were imaged during regeneration, and blastema size was quantified. Animals received an intraperitoneal injection of 5-ethynyl-2′-deoxyuridine (EdU) at 2 mg/kg one day before tissue collection.

### Imaging and quantification

Imaging was performed on Nikon Spinning Disk Confocal (CSU-W1), Zeiss Axio Scan.Z1, and a Leica M165 FC equipped with Leica DFC310 FX camera. Image quantification was conducted blindly. Cell counts were conducted using CellProfiler (*51*). Blastema area and HCR signal quantifications were measured using Fiji (*52*).

### Cell dissociation and single-cell RNA-sequencing

After hypophysectomy or sham treatment, blastemas, from 9 animals in each condition, were collected at 7 dpa. Each pooled replicate was dissociated in 1mL digestion buffer (100 µg/mL Liberase TM (Roche) diluted in 0.7X PBS with 0.5 U/µL DNase I (Invitrogen)) for 30 minutes at room temperature on a rocker. Suspensions were filtered through 40 µm strainers to remove cell debris and clumps. Strainers were washed with 1ml 0.7X PBS. Cells were spun down at 300xg for 5 minutes at room temperature. Cells were resuspended in 60 μL of 0.7X PBS-0.04% BSA. Tubes were delivered to Bauer Core Facility at Harvard University.

#### Single-cell RNA-sequencing

Each library pooled eight blastemas from two animals. Five dissociations were made per condition (HPS, sham) and the four with the highest cell count and viability were submitted to the Harvard University Bauer Sequencing Core for library preparation, yielding eight libraries total from sixteen animals. The unit of replication is the library (n=8, four per condition). The two animals within a library are pooled into a single measurement. Libraries were prepared with the Chromium GEM-X Universal 3’ Gene Expression v4 kit (10x Genomics) at a target recovery of 20,000 cells each and sequenced across three lanes of a NovaSeq X 25B flow cell (Illumina) with 2 x 151 cycles; reads were trimmed to 28bp and 90bp at demultiplexing.

Reads were pseudoaligned and collapsed to UMI-corrected gene counts via kb-python v0.29.5 (*53*), using the kallisto/bustools workflow and 10xv4 chemistry, with all lanes of a library quantified together. A reference index was generated via kb ref using the *Ambystoma mexicanum* UKY_AmexF1_1 genome assembly and RefSeq annotation (GCF_040938575.1), comprising 152,494 transcript sequences across 122,775 gene loci.

Cell calling, quality control and clustering used R 4.4.2 and Seurat v5.2.1 (*54*), while downstream analyses used R 4.3.3 and Seurat v5.0.3. Cells were called with emptyDrops (DropletUtils v1.26.0; (*55*)) at FDR 0.001 and a 500-UMI minimum. Ambient RNA was estimated with SoupX v1.6.2 (*56*). Corrected counts were used only for normalization and clustering, while differential expression and log-normalized expression use the uncorrected counts. Cells with fewer than 300 detected genes or in the lowest 3% of UMI counts in their library were removed, and doublets were removed with scDblFinder v1.20.2 (*57*) at an expected doublet rate of 20%. Libraries were merged without batch integration, and cells with ≥ 5% mitochondrial content were excluded. Counts were normalized using SCTransform v0.4.1 (*58*), with glmGamPoi v1.18.0 (*59*), regressing out mitochondrial fraction. Cells were clustered on the PCA nearest-neighbor graph and embedded by UMAP (dims 1-20, resolution 0.15, seed 5555), resulting in 15 manually annotated cell types across 147,253 cells. Where single-cell expression is shown directly, it is log-normalized expression, log1p of counts per 10,000. Epithelial cells and fibroblasts were subclustered separately through a similar workflow (SCTransform, 3,000 variable features, 20 principal components, resolution 0.4). Subcluster stability was confirmed by reclustering 100 random subsamples of 80% of cells (median Jaccard index 0.75-0.94 for the four subclusters analyzed)(*60*).

Differential expression was tested in the 13 cell types that were also subclustered (146,092 cells). Mast cells (909 cells) and granulocytes/monocytes (252 cells) were not subclustered and were not tested. Differential expression was assessed on pseudobulk profiles, by summing raw counts across cells of each library within cell type (libraries with ≥10 cells of that type; genes with ≥10 counts in at least two libraries) and testing them in DESeq2 v1.42.1 (*61*) by Wald test, applying Benjamini-Hochberg correction. The same pipeline was repeated at subcluster level, with correction within each subcluster, restricted to subclusters contributing at least two libraries per condition, each contributing ten or more cells. Volcano plots show ashr-shrunken log2 fold changes (*62*). All adjusted p-values come from the unshrunk maximum-likelihood fit.

The UKY_AmexF1_1 annotation assigns a gene symbol to 69,411 of 122,775 loci. For genes reported here that lacked a symbol, gene identity was established by reciprocal best-hit BLAST together with the RefSeq product description, and the LOC identifier is reported alongside the resolved symbol. Loci that could not be resolved retain their LOC identifier alone.

#### Pituitary Bulk RNA-sequencing

Animals measuring 8-12cm were amputated unilaterally at the stylopod, with 15 animals harvested per timepoint. Pituitary glands were collected from intact axolotl salamanders at 0, 1, 3, 7, and 14 days after limb amputation. Animals were perfused before collection, and every animal received an EdU injection the day before harvest. Three sequencing samples were prepared per timepoint, 15 samples in total, each pooling glands from five animals. RNA was extracted using the NucleoSpin RNA Plus XS kit (Macherey-Nagel). Libraries were prepared by the Harvard University Bauer Sequencing Core using the Watchmaker RNA Library Prep Kit (Watchmaker Genomics) with poly-A selection, and sequenced on a NovaSeq X 10B flow cell (Illumina) as 151bp paired-end reads to 84-116 million read pairs per sample.

Transcript abundances were estimated with kallisto v0.46.0 (*63*) against an index built from the RefSeq transcript set of the UKY_AmexF1_1 assembly (GCF_040938575.1; 96,841 transcripts) and summarized to gene level with tximport v1.30.0 (*64*). Counts were analyzed in DESeq2 v1.42.1, retaining 19,323 genes after filtering. An overall effect of time was tested by likelihood-ratio test against the intercept-only model, and each timepoint was compared with intact by Wald test, both with Benjamini-Hochberg correction. Transcript abundance shares are computed from kallisto TPM. Gene sets were tested with limma fry and camera on voom-transformed counts; sets with padj < 0.05 on both sets, in the same direction, are reported.

#### Mouse digit tip scRNA-seq analysis

Processed single-cell RNA sequencing data from regenerating P3 digit tips were provided by the Lehoczky laboratory as the annotated Seurat object described in (*28*), in which reads were counted with Cell Ranger 7.1 including intronic reads. These datasets are publicly available: unamputated (UA) and 11, 12, 14, 17 dpa from(*29*), and 28 dpa from (*30*) (GEO GSE143888 and GSE267446). We subsetted the object to the 43,408 P3 cells (one library per timepoint) and retained the published cell type labels. Seurat v5.0.3 in R 4.3.3 was used for analysis. Counts were log-normalized and data was scaled and reduced by principal component analysis (30 PCs) without integration and used for UMAP construction. Fibroblasts were reclustered using the same methods (20 components; res 0.6) into 16 subclusters, annotated using FindAllMarkers (Wilcoxon rank-sum test; min.pct 0.25; log2 fold change >0.25) as described(*28*). Dot plots show percentage of cells within each group with ≥1 UMI and mean log-normalized expression per group, pooled across timepoints.

For *Prlr+* versus *Prlr-*comparisons, each *Prlr+* fibroblast (≥1 *Prlr* UMI; 11-28dpa; n=86) was matched to up to 10 *Prlr*-fibroblasts from the same subcluster, timepoint, and sequencing-depth quintile (n = 847). Genes were compared using FindMarkers (Wilcoxon rank-sum test) and identified as significant (padj < 0.05; detected in ≥ 25% of cells in either group; |log2 fold change| > 0.58 following (*30*)).

#### Bulk RNA-seq analysis of mouse P3 nail epithelium

Hindlimb digits 2-5 of 6-week-old female FVB/NJ mice were P3 amputated and collected from unamputated digits and at 1, 3, 6, 9 and 12 dpa. Each sample pooled 16 digits from 2 mice (n = 3 samples per time point). Nail epithelium was microdissected and dissociated with trypsin and collagenase, and 1,500 *CD49f*+ cells per sample (anti-CD49f–APC, BioLegend 313609; 1:200) were sorted directly into TRIzol. RNA was used for ultra-low-input, strand-specific library construction and sequencing (Genewiz). Reads were trimmed with Trimmomatic v0.36, aligned to the GRCm38 genome with STAR v2.5.2b, and counted over Ensembl (release 87) exons with featureCounts (Subread v1.5.2), using uniquely mapped, strand-specific reads. For each time point versus unamputated, genes with a mean raw count above 10 across the six samples were tested in DESeq2 v1.18 (Wald test, Benjamini-Hochberg correction), and maximum-likelihood log2 fold changes are reported. We re-ran this analysis from the deposited count files and reproduced the original results. Data is available from GEO Accesssion GSE219177.

## Acknowledgments

We acknowledge Christopher Morii-Sciolla for assistance with sorting and sequencing mouse P3 nail epithelium samples.

This material is based upon work supported by the National Science Foundation Graduate Research Fellowship under Grant No (2140743). Any opinion, findings, and conclusions or recommendations expressed in this material are those of the authors(s) and do not necessarily reflect the views of the National Science Foundation.

We acknowledge the use of Claude (Anthropic) to help write analysis code for the transcriptomic datasets and to edit the manuscript for grammar and readability. The authors reviewed and verified all AI-assisted code and text and take full responsibility for the content of the work.

## Funding

NSF CAREER IOS-2145925

NICHD 1R01HD115272

Harvard University Faculty of Arts and Sciences Dean’s Competitive Fund HCBI Simmons Award

NIH/NICHD R01HD109200

AFSH Basic Science Award #1399

## Author contributions

Conceptualization: NL, JCP, DP, APS, JPB, JLW

Methodology: NL, JCP, DP, HS, MR, JAL, APS, JPB, JLW

Investigation: NL, JCP, DP, HS, TF, LLU, AZ, ER, ARM, BKF, RM, RC, TG, SYCW, LS, AJ, EC, AES, AL, RTK, MXT, ATH, TSP, MC, ASG, APS, JPB

Visualization: JCP, NL, DP, HS

Funding acquisition: JAL, APS, JPB, JLW

Supervision: APS, JLW

Writing – original draft: JCP, NL, HS, JLW

Writing – review & editing: all authors

## Data, code, and materials availability

Single-cell RNA-seq data (axolotl HPS and sham blastemas) and bulk RNA-seq data (axolotl pituitary) have been submitted to GEO and will be made public on journal publication. The mouse P3 digit single-cell libraries are available under GEO GSE143888 and GSE267446 (*29, 30*), and the annotated count object used here is available from the Lehoczky laboratory on request. The mouse nail epithelium bulk RNA-seq data were generated by the Lehoczky laboratory and will be released by them on publication. Analysis code will be made available on publication. All other data are available in the main text or the supplementary materials.

---

**(S1).**
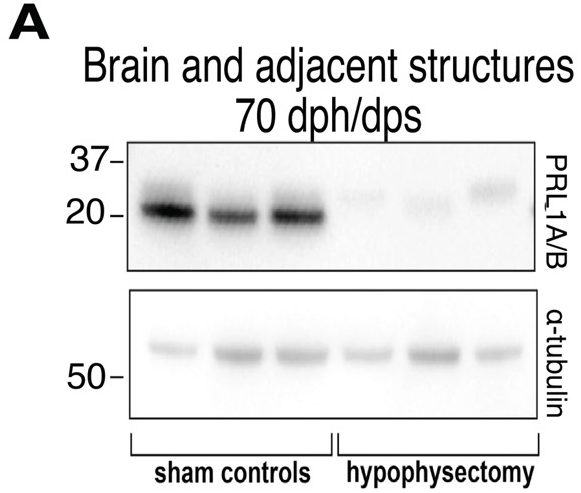
Western blot showing pituitary does not regenerate up to 70 days post hypophysectomy. **(A)** Six lanes: 3 brains and adjacent tissues from sham-operated animals and 3 brains and adjacent tissues from HPS-operated animals. Probed for PRL1 and alpha tubulin as a loading control.

**(S2).**
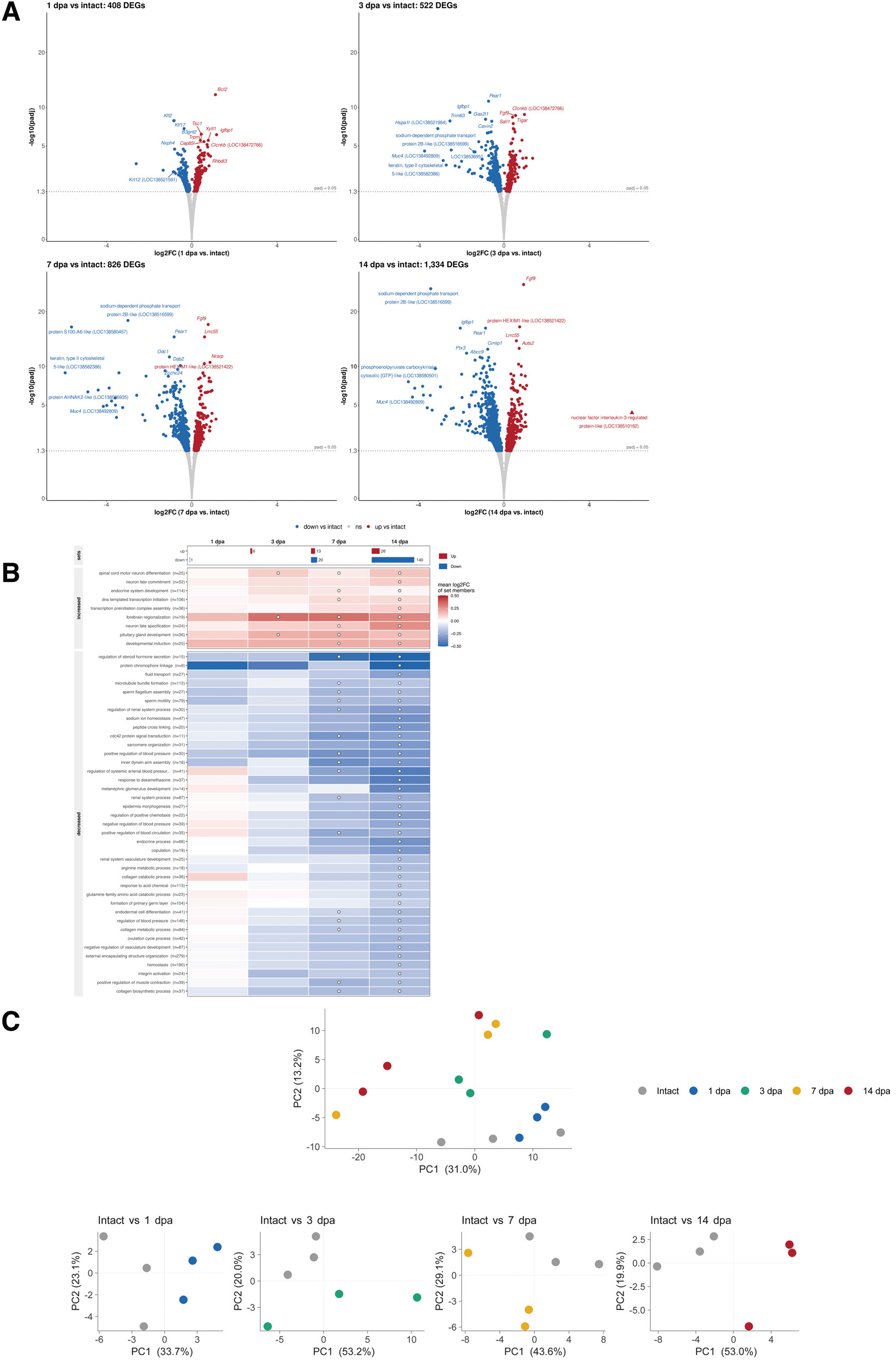
Limb amputation results in a coordinated transcriptional response in the pituitary gland, accumulating over time. **(A)** Volcano plots of differentially expressed genes in pituitary at each timepoint after limb amputation compared to intact. **(B)** Gene sets significantly altered in the pituitary gland at 1, 3, 7, and 14 dpa versus intact, by direction (GO biological process; 4,220 sets tested per timepoint; rotation testing with limma *fry*, restricted to sets also significant under *camera* in the same direction, BH < 0.05). Significant sets accumulate over the time course and are visualized by bars at the top of the figure (1dpa = 1 set, 3dpa = 6 sets, 7dpa = 33 sets, and 14dpa = 166 sets). Mean log2 fold change is visualized through cell color; white dots mark significance. **(C)** Principal components for all 15 bulk pituitary libraries (intact, 1, 3, 7, and 14 dpa; n = 3 per timepoint; PC1 31.0%, PC2 13.2% variance). The smaller panels are recomputed on intact versus a single timepoint. PC1 carries 33.7%, 53.2%, 43.6% and 53.0% of variance for 1, 3, 7 and 14 dpa respectively, and separates the intact condition from amputated glands at every timepoint.

**(S3).**
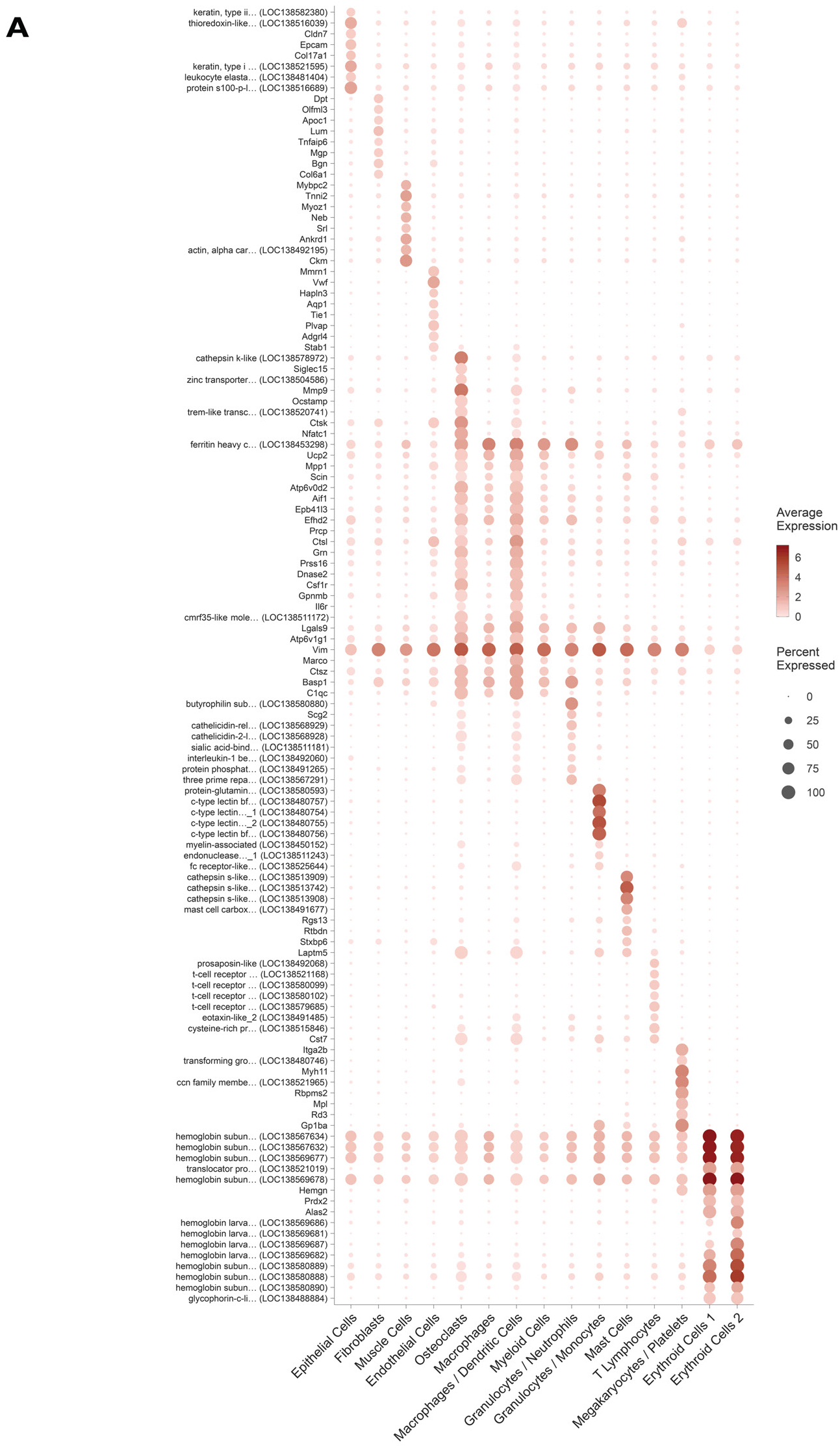
Marker genes for the 15 cell types in the hypophysectomy vs. sham scRNA-seq dataset. **(A)** Dot plot of the marker genes used to annotate the 15 cell types in the HPS vs. Sham dataset (dot size = % of cells with ≥1 UMI; color = mean log-normalized expression within cell type). Eight markers are shown per cell type. Genes are ranked within each cell type by most selectively enriched. Color is absolute expression, so values are comparable across genes. Only genes with existing NCBI annotations are shown, and LOC-identification only genes were dropped. Mitochondrial transcripts were excluded from marker selection – they ranked highest for Myeloid Cells but are expressed across every cell type and reflect possible cell stress rather than identity.

**(S4).**
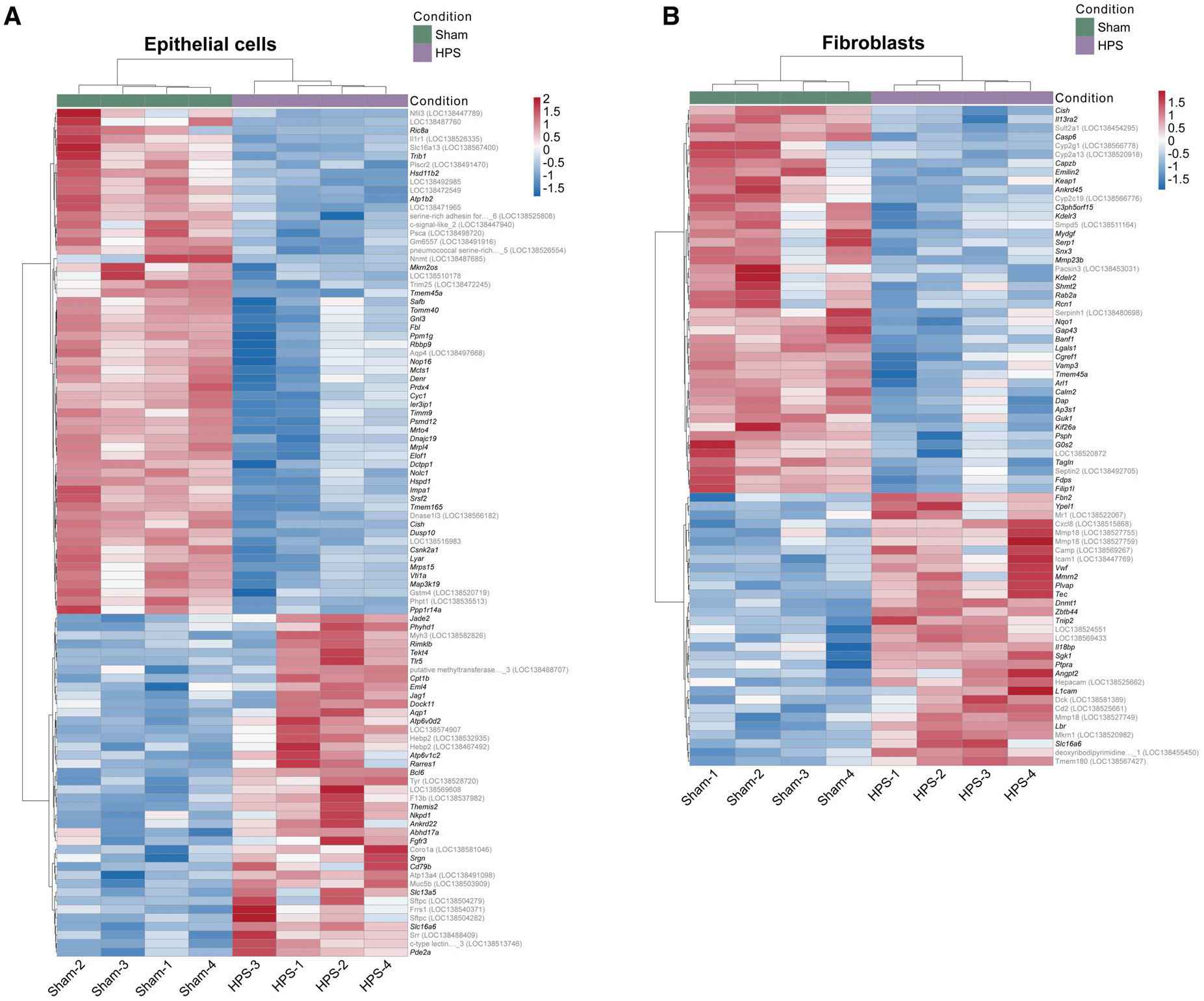
Epithelial genes that change after hypophysectomy are consistent across all eight libraries. Fibroblast genes that change after hypophysectomy are consistent across all eight libraries. **(A)** Per-library pseudobulk expression heatmap of the 99 genes significant in Epithelial Cells (padj < 0.05, HPS vs. Sham), standardized within each gene (row z-score) and clustered on both axes (ward.D2). Each column is one library (4 Sham, 4 HPS), and the bar across the top marks surgical condition (Sham = green; HPS = purple). Gene symbols are italicized; grey rows are loci lacking a gene symbol in the NCBI annotation. Each library follows its condition and no single library drives its signal. Row z-scores are within each gene and are not comparable across genes. **(B)** Per-library pseudobulk expression heatmap of the 75 genes significant in Fibroblasts (padj < 0.05, HPS vs. Sham), standardized within each gene (row z-score) and clustered on both axes (ward.D2). Each column is one library (4 Sham, 4 HPS), and the bar across the top marks surgical condition (Sham = green; HPS = purple). Gene symbols are italicized; grey rows are loci lacking a gene symbol in the NCBI annotation. Each library follows its condition and no single library drives its signal. Row z-scores are within-gene and are not comparable between genes.

**(S5).**
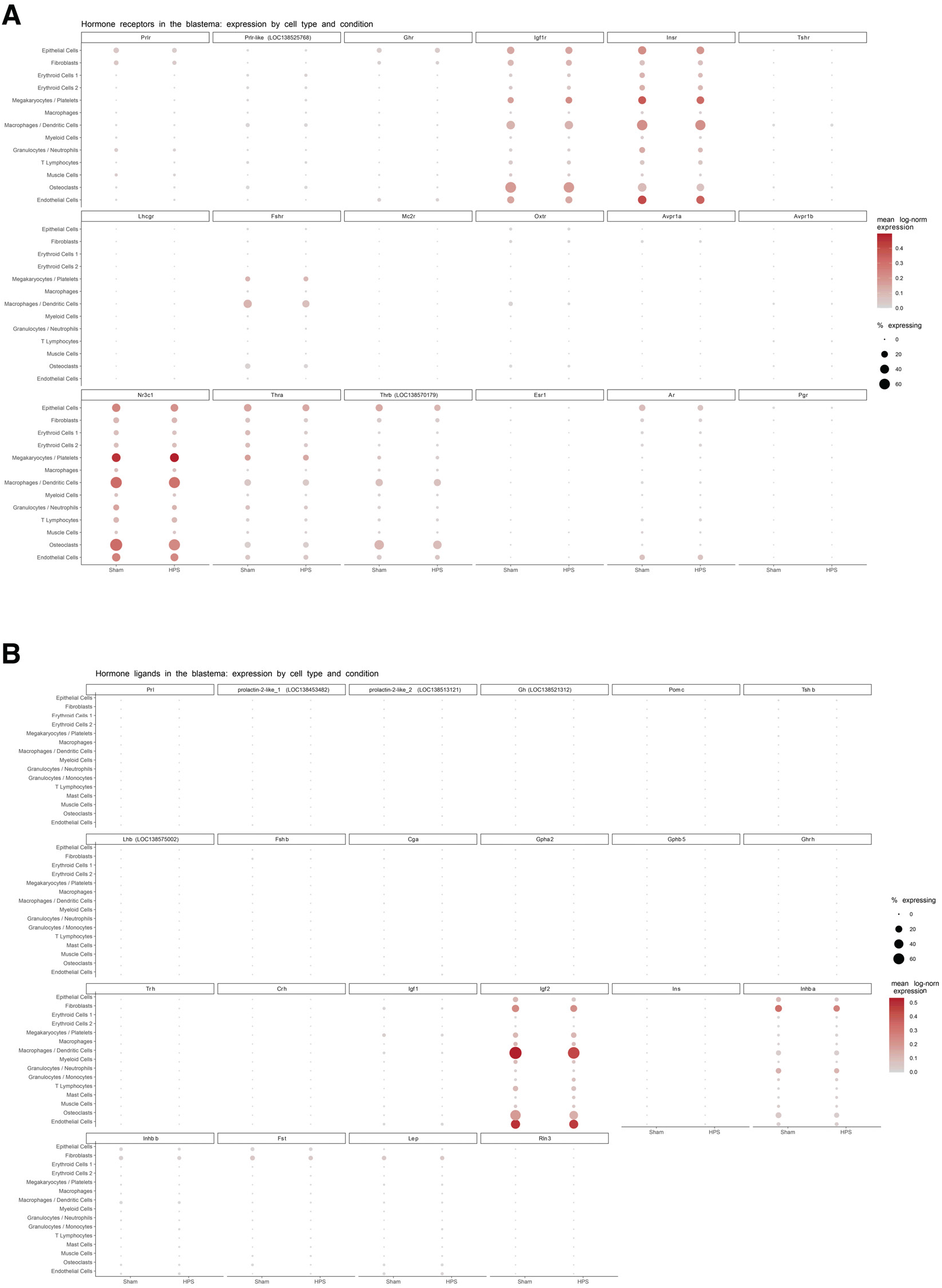
Canonical pituitary hormone receptor abundances do not change after hypophysectomy in any cell type. Canonical pituitary hormone ligands are largely absent from the blastema. **(A)** Dot plot of 18 hormone receptors by cell type and surgical condition (dot size = percentage of cells with ≥1 UMI; color = mean log-normalized expression; Sham and HPS shown side-by-side within each cell type). No receptor reaches significance in any cell type via pseudobulk differential expression (padj < 0.05; DESeq2 Wald, BH-corrected within cell type). **(B)** Dot plot of 22 hormone ligands by cell type and surgical conditions (dot size = percentage of cells with ≥1 UMI; color = mean log-normalized expression; Sham and HPS shown side-by-side within each cell type). No ligand reaches significance in any cell type via pseudobulk differential expression (padj <0.05; DESeq2 Wald, BH-corrected within cell type).

**(S6).**
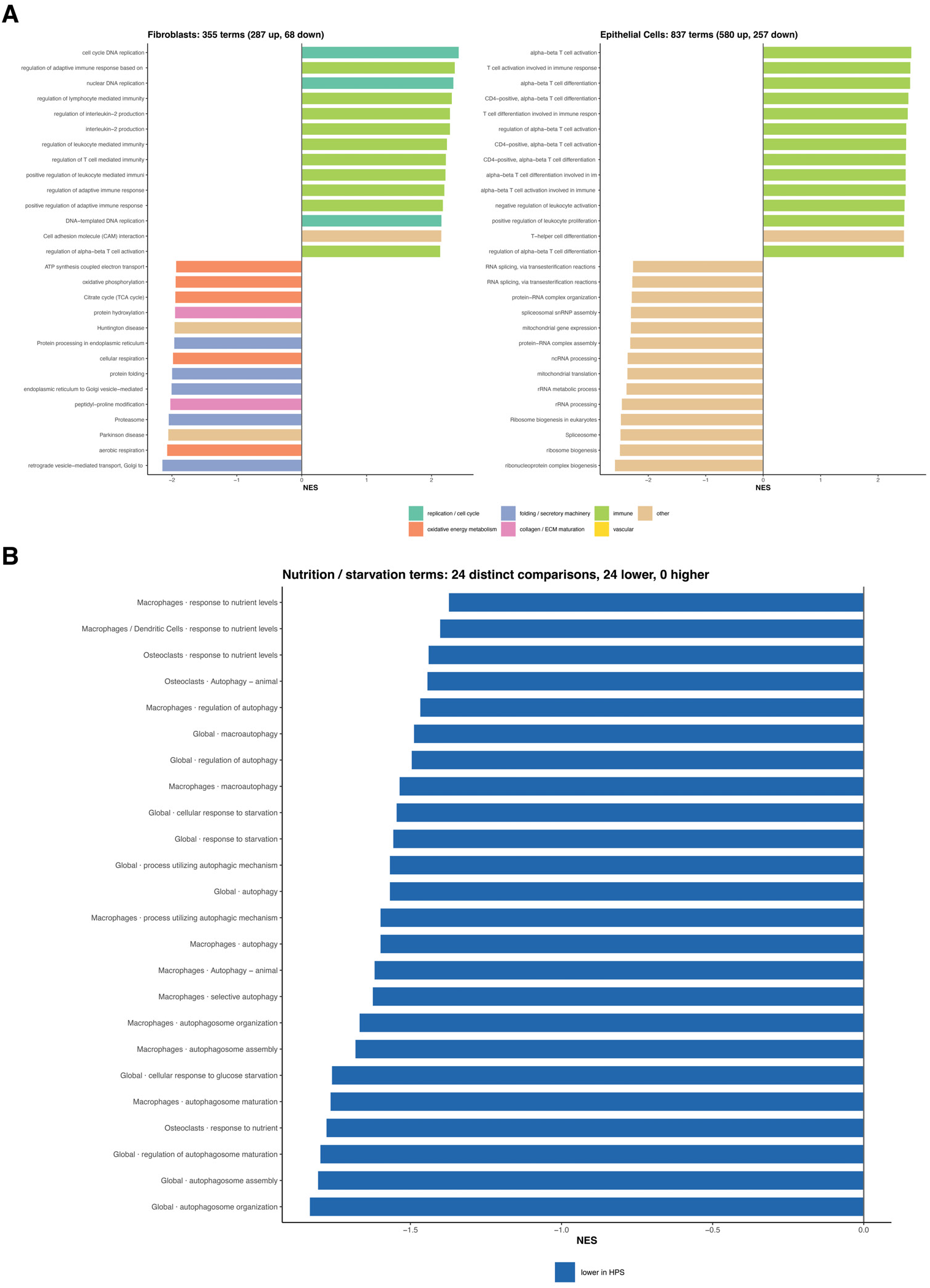
Hypophysectomy affects fibroblasts and epithelial cells in different ways. **(A)** Gene set enrichment analysis (GSEA) of fibroblasts and epithelial cells, HPS vs. Sham (GO biological process and KEGG pathways), showing top terms by normalized enrichment score (NES) in each direction. In fibroblasts, 355 terms move (287 up; 68 down); all 11 DNA-replication terms rise, while respiration, protein folding, and proteasome fall. In epithelial cells, 837 terms move (580 up; 257 down), with lymphocyte-activation programs up and RNA-processing programs down. **(B)** Starvation, autophagy and nutrition gene sets by cell type, HPS vs. Sham. Of 24 significant comparisons, all 24 are lower in HPS (17/17 autophagy; 3/3 starvation; 4/4 nutrient gene sets). All GSEA terms shown are significantly downregulated in HPS condition, opposite of what would be expected if HPS animals were malnourished.

**(S7).**
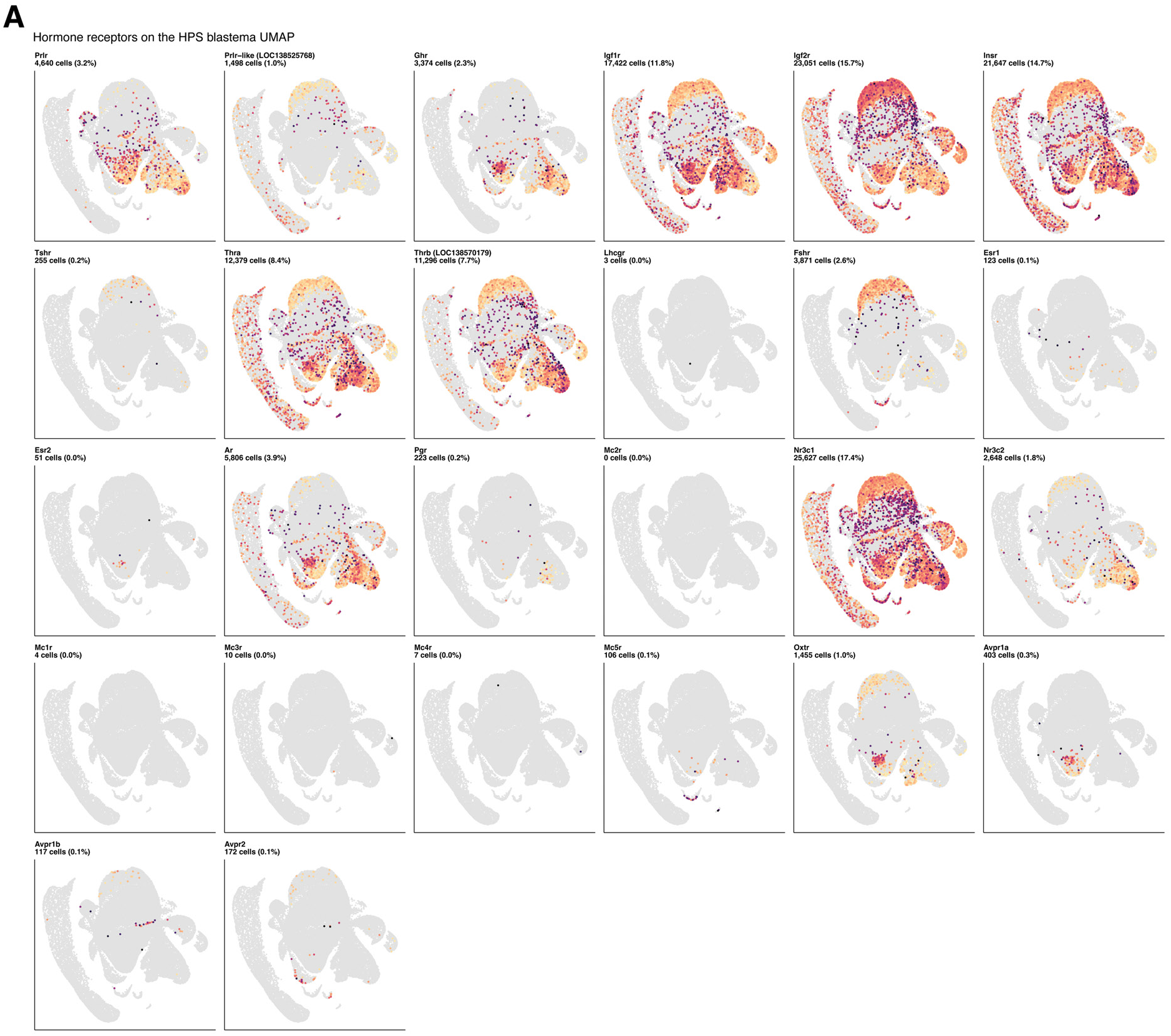
Pituitary hormone receptor expression varies broadly across the blastema. **(A)** UMAP feature plots for 26 hormone receptors across the full HPS vs. Sham dataset (147,253 cells; 15 cell types), grouped by endocrine axis. Grey = no UMI detected; color = log-normalized expression, with each panel scaled independently so presence and pattern are readable but intensity is not comparable across panels.

**(S8).**
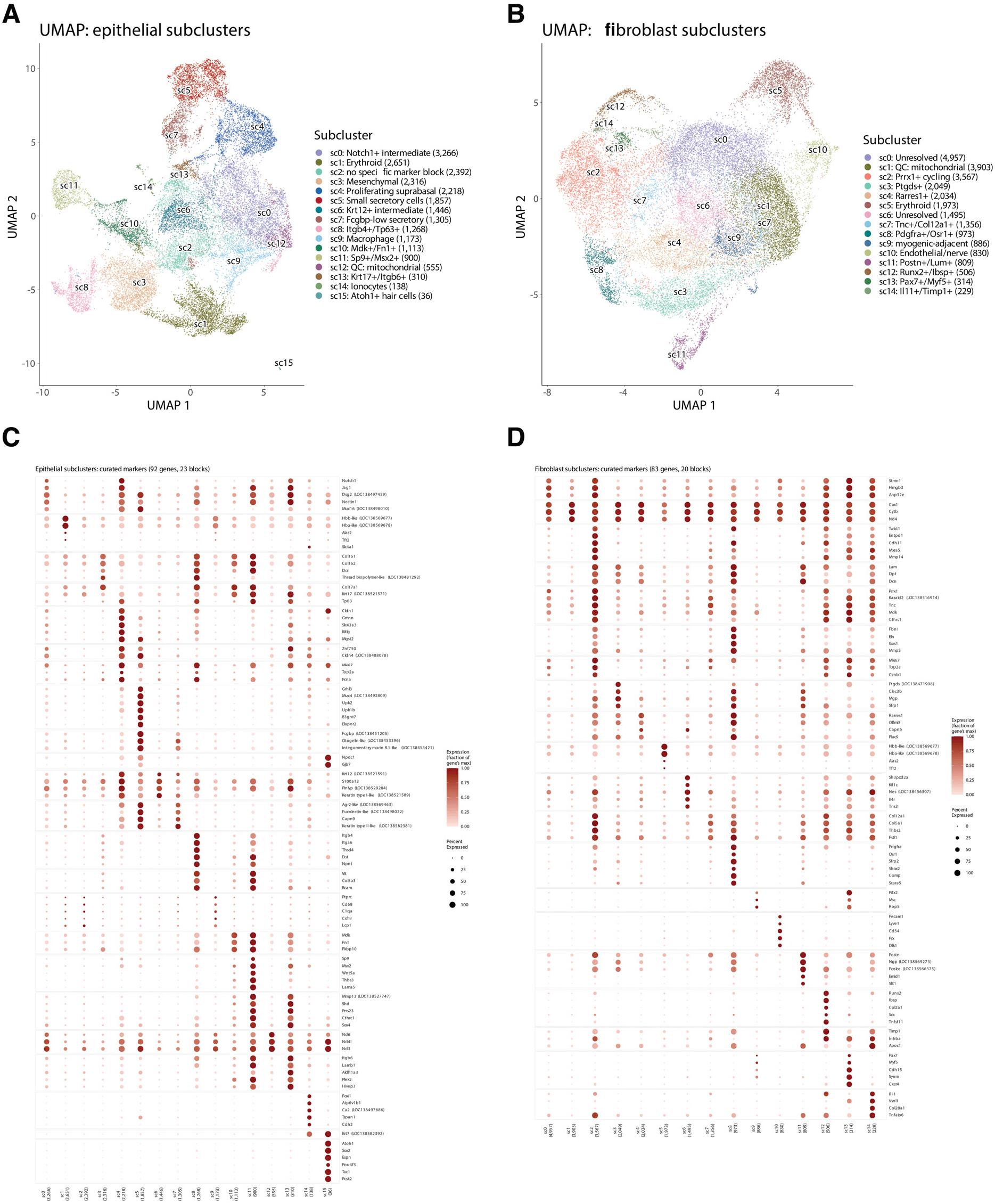
Reclustering of epithelial cells and fibroblasts reveals cell subtype diversity in pituitary-responsive cells. **(A)** UMAP of the 22,944 epithelial cells reclustered into 16 subclusters. **(B)** UMAP of 25,881 fibroblasts reclustered into 15 subclusters. **(C)** Dot plot of curated marker genes of the 16 epithelial subclusters, all cells (Sham and HPS). Dot size = % of cells with ≥1 UMI; color = mean log-normalized expression scaled to each gene’s maximum across subclusters. **(D)** As in (C), for the 15 fibroblast subclusters.

**(S9).**
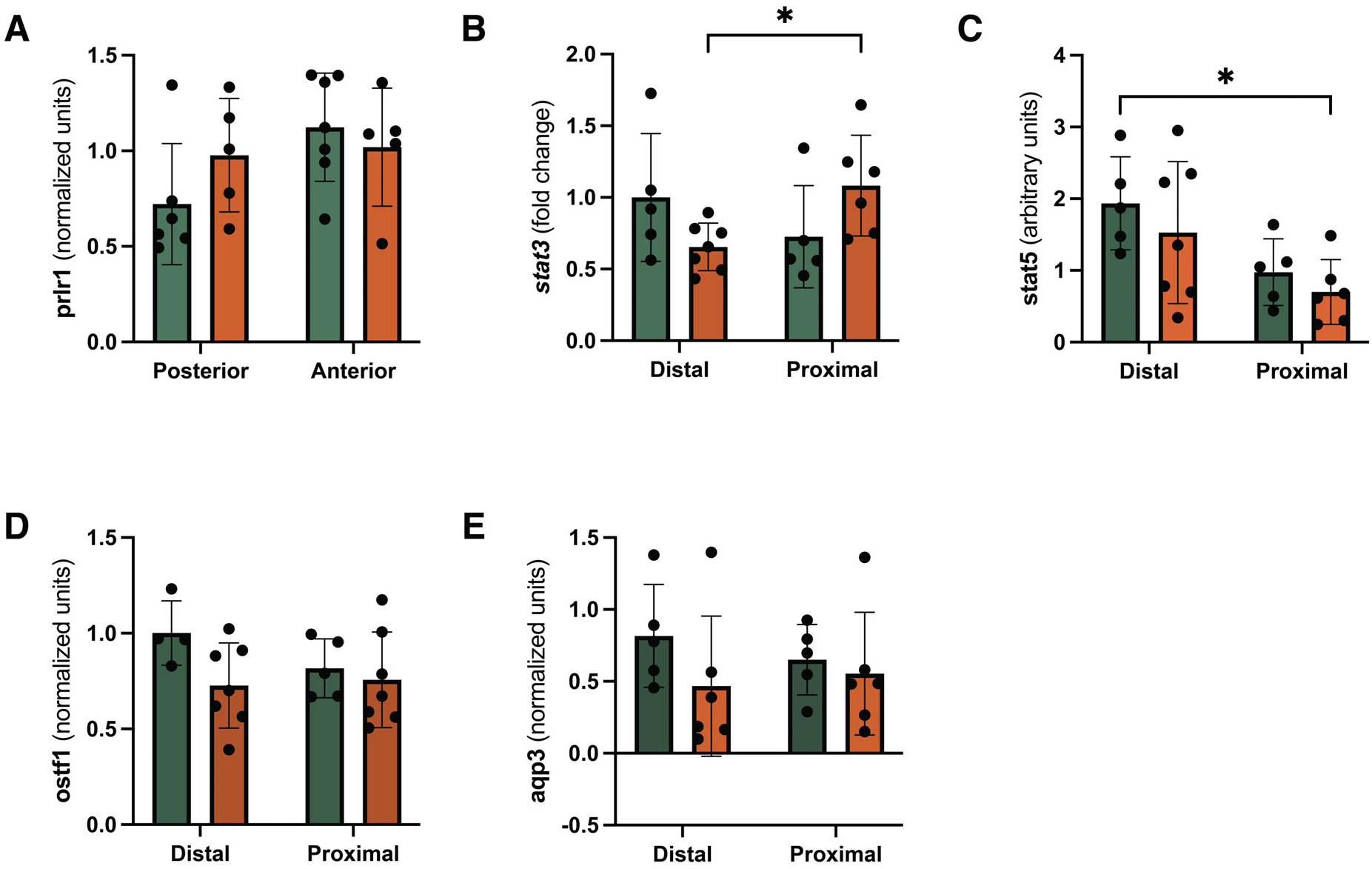
qPCR fold-change for endocrine related pathways. **(A)** *Prlr1* (qPCR), same design as Fig. 4D. Two-way ANOVA and Tukey’s multiple comparisons test. **(B)** *Stat3* (qPCR), same design as Fig. 4D. Two-way ANOVA and Tukey’s multiple comparisons test. *: p < 0.05 **(C)** *Stat5* (qPCR), same design as Fig. 4D. Two-way ANOVA and Tukey’s multiple comparisons test. *: p < 0.05 **(D)** *Ostf1* (qPCR), same design as Fig. 4D. Two-way ANOVA and Tukey’s multiple comparisons test. **(E)** *Aqp3* (qPCR), same design as Fig. 4D. Two-way ANOVA and Tukey’s multiple comparisons test.

**(S10).**
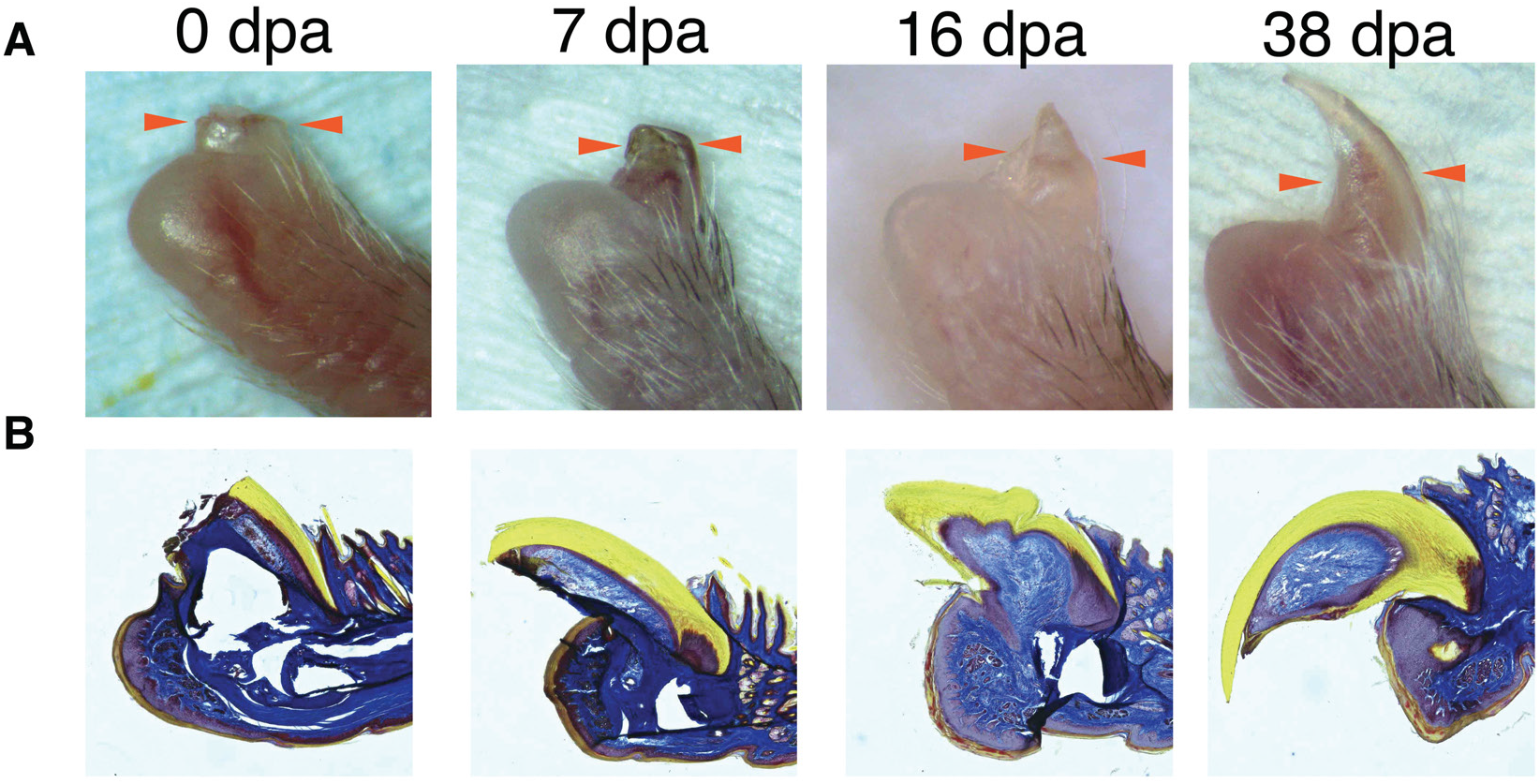
Morphology and histology of P3 digit tip regeneration. **(A)** Representative images of the mouse P3 digit regeneration time course (0, 7, 16, 38 dpa); orange arrowheads mark the amputation/regenerating tip.**(B)** Masson’s trichrome staining of the same P3 digit regeneration time course (0, 7, 16, 38 dpa).

**(S11).**
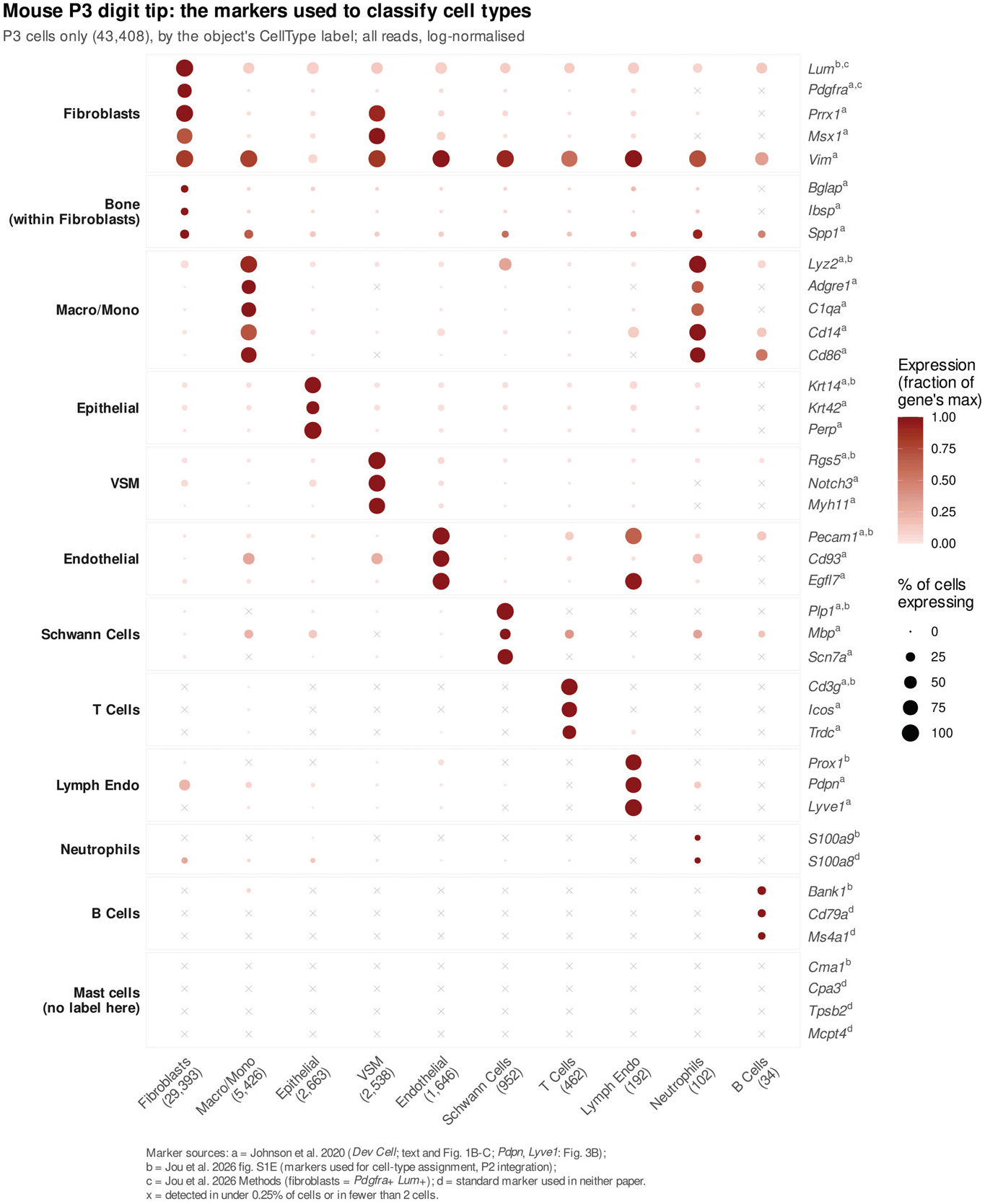
Marker genes for the 10 published cell types in the P3 mouse digit amputation scRNA-seq dataset.

**(S12).**
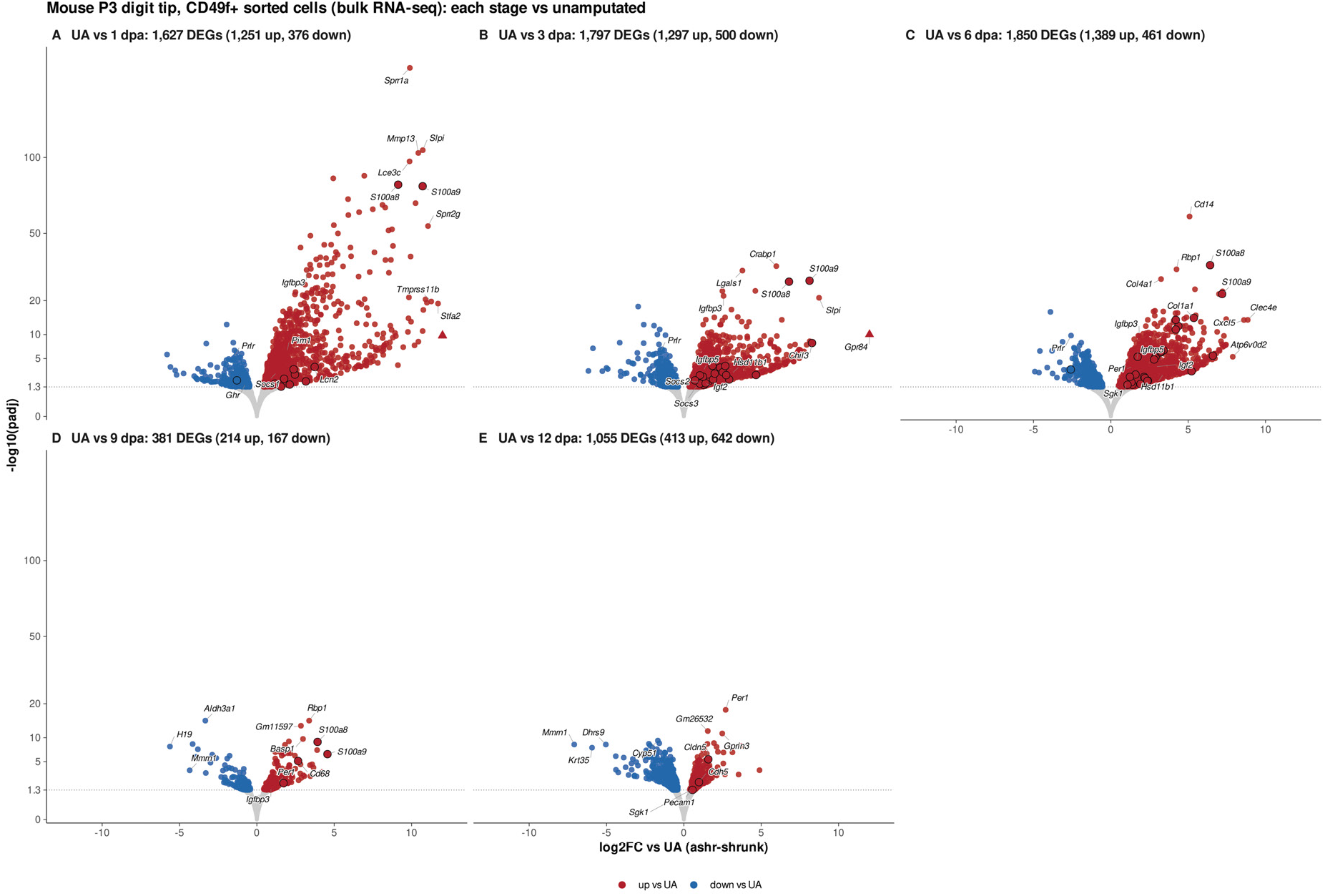
Volcano plots of DEGs for each amputation timepoint compared to unamputated for *CD49f*+ sorted nail epithelium.

